# Glycolysis-derived alanine from glia fuels neuronal mitochondria for memory in Drosophila

**DOI:** 10.1101/2022.05.30.493961

**Authors:** Yasmine Rabah, Raquel Frances, Pierre-Yves Plaçais, Thomas Preat

**Author notes:** co-senior authors. Correspondence,.

## Abstract

Glucose is the primary source of energy for the brain. However, it remains controversial whether, upon neuronal activation, glucose is primarily used by neurons for ATP production, or if it is partially oxidized in astrocytes, as proposed by the astrocyte-neuron lactate shuttle model for glutamatergic neurons. Thus, an *in vivo* picture of glucose metabolism during cognitive processes is missing. Here, we uncover in Drosophila a glia-to-neuron alanine transfer that sustains memory formation. Following associative conditioning, glycolysis in glial cells produces alanine, which is back-converted into pyruvate in mushroom body cholinergic neurons to uphold their increased mitochondrial needs. Alanine, as a mediator of glia-neuron coupling, could be an alternative to lactate in cholinergic systems. In parallel, a dedicated glial glucose transporter imports glucose specifically for long-term memory, by directly transferring it to neurons for use by the pentose phosphate pathway. Our results demonstrate *in vivo* the compartmentalization of glucose metabolism between neurons and glial cells during memory formation.

## Introduction

Glucose is the primary energy metabolite consumed by the brain^1^, for which neuronal electrical activity and neurotransmission are the major energy sinks. Glucose is transported from the blood circulation into both neurons and neighboring glial cells, particularly astrocytes^2^. However, determining which of the two cell types is the primary site of glucose uptake accompanying increased neuronal activity has long been a matter of controversy^3,4^. The astrocyte-neuron lactate shuttle (ANLS) is one hypothesized synaptic mechanism in which the astrocytes are the main glucose consumers. According to this model, astrocytes transfer glycolysis-derived lactate to neurons upon glutamate reuptake^5,6^. In turn, lactate can be used by neurons for signaling^7^ or oxidative purposes and energy production. The observations that astrocytes can be net suppliers of lactate and that the glycolysis rate is higher in astrocytes than in neurons are well-supported^4^. Several studies have shown that long-term memory (LTM) in particular relies on lactate transfer from astrocytes to neurons^8–10^. However, other studies continue to challenge the ANLS scheme, as they have shown that glucose is preferentially consumed by neurons upon acute stimulation^11,12^. In support of this opposing model, it was reported that neuronal glycolysis is upregulated for energy production in response to neuronal stimulation^13^. Therefore, although the ANLS model aptly solves how energy supply is tailored to synaptic activity, its relevance *in vivo* is still disputed^3,14^. Moreover, the precise fate of glucose in upholding neuronal oxidative metabolism *in vivo*, especially during complex cognitive functions like memory, remains elusive. Notably, the controversy regarding the compartmentalization of glucose metabolism between neurons and glia has crystallized around the ability of lactate to fuel neuronal oxidative metabolism, leaving alternative metabolic derivations of glial glycolysis largely under-investigated. For instance, the L-amino acid alanine is derived from glial glucose metabolism and can serve as a metabolic substrate^15–17^, making it a potential substitute for lactate.

The unmatched genetic, behavioral, and cellular-resolution imaging toolsets available to Drosophila studies provide a powerful and temporally precise means to investigate, *in vivo*, the metabolic processes engaged in high-order brain functions like memory. Drosophila can form aversive olfactory memory upon the association between an odorant and electric shocks, and a variety of conditioning protocols have been designed that evoke memories with different properties and persistence^18,19^. All forms of olfactory memories are encoded within specific subsets of neurons in the mushroom body (MB), a bilateral structure of about 2,000 cholinergic neurons in each brain hemisphere^20^. Drosophila neurons are surrounded by several types of glial cells, making this model relevant for the study of neuron-glia metabolic coupling supporting memory formation.

Recently, we successfully unveiled several metabolic adaptations *in vivo* that underlie memory formation, including neuron-glia metabolic coupling processes^21,22^. We found that the early phase of LTM formation is sustained by the rapid upregulation of neuronal pyruvate metabolism by mitochondria^23^. Furthermore, we revealed the existence of a concomitant glucose shuttle from perisomatic glial cells (known as cortex glia) to neuronal somata that supplies the neuronal pentose phosphate pathway (PPP) specifically for LTM formation, such that MB neuron glucose utilization is uncoupled from pyruvate oxidation for LTM^21^. Because the cellular and molecular origins of pyruvate sustaining memory had not yet been determined, this prompted us to identify the mechanisms supporting upregulated pyruvate metabolism for memory formation.

Here, we have uncovered the reliance of specific forms of aversive memories, i.e. middle-term memory (MTM) and LTM, on pyruvate metabolism in MB neurons. Our results indicate that the source of neuronal pyruvate for both MTM and LTM is the amino acid L-alanine. We further show that alanine is produced by neighboring cortex glia through glycolysis. Finally, we revealed that two distinct glucose transporters contribute in parallel to glucose import by glial cells for distinct fates: one for glial glycolysis that sustains MTM and LTM, and one for direct glucose transfer to MB neurons specifically for LTM. Overall, our results demonstrate the *in vivo* compartmentalization of glucose metabolism between neurons and glial cells for memory formation, with alanine transfer from glia to neurons bridging glial glycolysis with neuronal mitochondrial pyruvate consumption.

## Results

### Mitochondrial pyruvate metabolism underlies the formation of specific memory phases

Following a single pairing of electric shocks with an odorant (single-cycle training), associative odor-avoidance memory persists for several hours but decays within 1 day. Based on genetic and circuit analyses, this memory was dissected into two distinct components, designated as middle-term memory (MTM)^18,19^ and middle-term anesthesia-resistant memory (MT-ARM). A stable protein synthesis-dependent LTM, which persists for several days, can be formed by subjecting flies to repeated conditioning cycles separated by rest intervals (5x spaced training)^18,19^. MTM and LTM are encoded in the same subset of MB neurons and retrieved through the same downstream circuits^24,25^, so that LTM formation is thought to rely on initial cellular substrates that are shared with MTM. Notably, repeated conditioning cycles without rest (5x massed training) induce a distinct form of consolidated memory (LT-ARM). This memory differs from LTM in terms of persistence^18^, the underlying neuronal circuits^25^ and support genes^26,27^, and metabolic regulations^21,23^.

Pyruvate import into mitochondria *via* the mitochondrial pyruvate carrier Mpc1^28^ is an obligatory step in the oxidative metabolism of glucose derivatives. In order to probe the reliance of Drosophila memory formation on neuronal pyruvate mitochondrial metabolism, we investigated the effect of impaired pyruvate uptake on memory using RNAi interference (RNAi) targeted against Mpc1 expression. For this, an Mpc1 RNAi was expressed selectively in MB neurons using the VT30559-GAL4 driver^23^. To avoid putative developmental issues, the thermosensitive GAL4 inhibitor GAL80^ts^ was ubiquitously coexpressed (tub-GAL80^ts 29^; see Methods section for details) so that GAL4-mediated RNAi expression would be induced only at the adult stage by heat activation for 3 days before conditioning. After single-cycle training, 3-hour memory was strongly but partially impaired (Fig. 1a). Cold shock treatment, which erases the labile MTM component while leaving the MT-ARM component intact, revealed that this impairment corresponds to a complete loss of MTM (Fig. 1a). LTM, measured 24 hours after spaced training, was also fully abolished by Mpc1 knockdown (Fig. 1a). By contrast, LT-ARM, measured 24 hours after massed training, was not affected by Mpc1 knockdown (Fig. 1a). This is in agreement with our previous report that LTM formation relies on the pyruvate dehydrogenase (PDH) mitochondrial enzyme complex and involves increased mitochondrial pyruvate consumption in MB neurons^23^. MTM and LTM performances were normal when RNAi expression was not induced (Extended Data Fig. 1a), which rules out the possibility that the observed memory defects could be due to leaky RNAi expression during development. We also verified that innate sensory acuity for the relevant stimuli was normal in the flies of interest (Extended Data Table 1). These behavioral experiments were replicated using a second, non-overlapping Mpc1 RNAi (Extended Data Fig. 1b). Together, these results demonstrate that MTM and LTM formation rely critically on mitochondrial pyruvate import into MB neurons.

**Figure 1:**
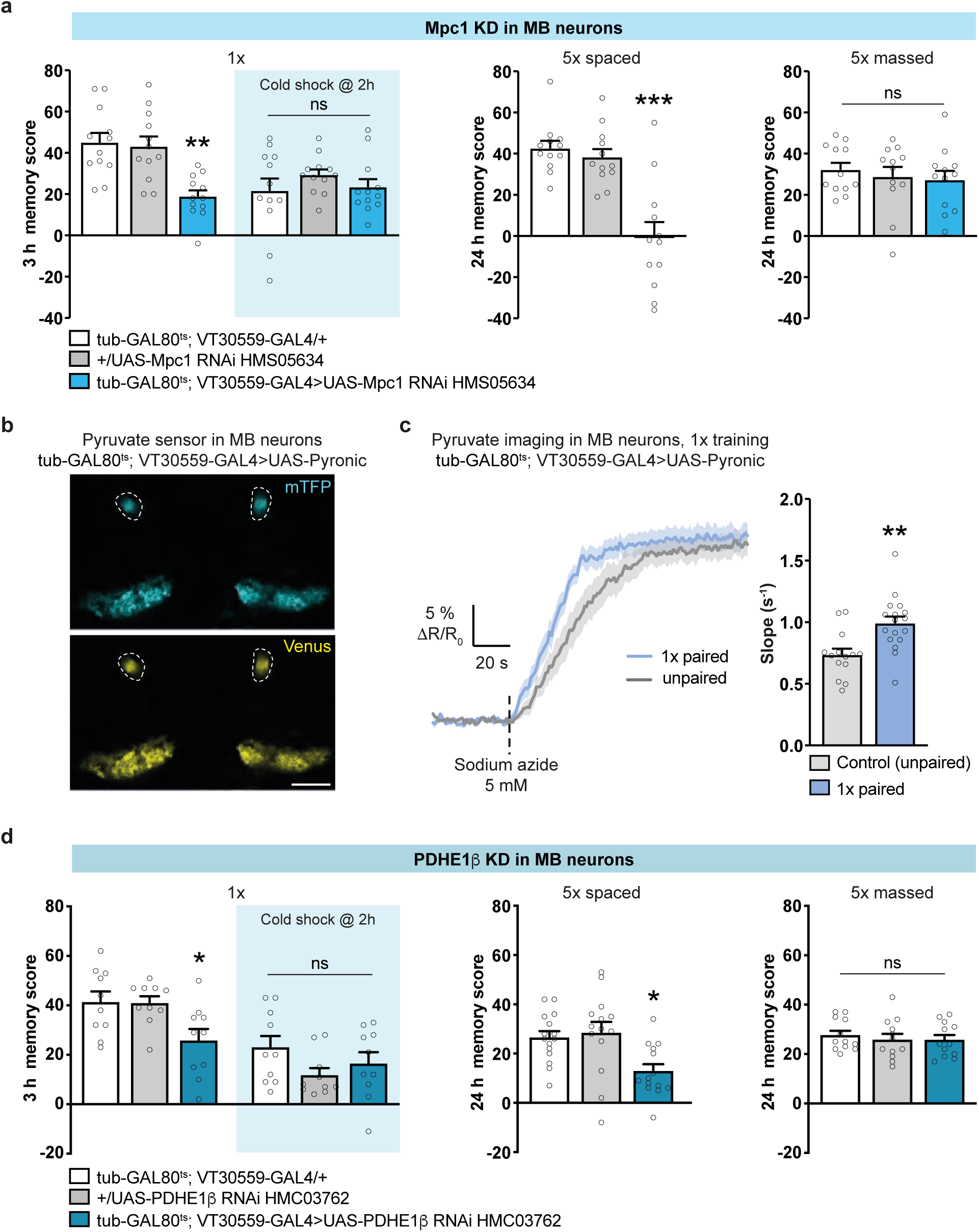
Pyruvate consumption by MB neurons increases upon memory formation. a. Mpc1 knockdown (KD) in adult MB neurons impaired memory after single-cycle (1x) training (n = 12, F_2,33_ = 11.40, P < 0.001) and spaced training (n = 12, F_2,33_ = 19.36, P < 0.001), but did not affect memory after single-cycle training followed by cold shock (n = 12, F_2,33_ = 0.83, P = 0.45) or massed training (n = 12, F_2,33_ = 0.36, P = 0.70). b. The pyruvate sensor Pyronic was expressed in adult MB neurons. The two images show the mTFP and Venus channels. The dashed lines delimit bundles of MB neuron axons, namely the vertical lobes, where the pyruvate FRET signal was quantified. Scale bar: 50 µm. c. Single-cycle training elicited a faster pyruvate accumulation in MB neuron axons following sodium azide application (5 mM) as compared to non-associative unpaired training (n = 14-16, t_28_ = 3.39, P = 0.002). d. PDHE1? knockdown in adult MB neurons impaired memory after single-cycle training (n = 10, F_2,27_ = 5.08, P = 0.01) and spaced training (n = 14-15, F_2,40_ = 6.25, P = 0.004), but did not affect memory after single-cycle training followed by cold shock (n = 10, F_2,27_ = 1.91, P = 0.17) or massed training (n = 12, F_2,33_ = 0.27, P = 0.76). All data are presented as mean ± SEM. Asterisks illustrate the significance level of the t-test, or of the least significant pairwise comparison following an ANOVA, with the following nomenclature: *p < 0.05; **p < 0.01; ***p < 0.001; ns: not significant, p > 0.05.

Using *in vivo* pyruvate imaging experiments to monitor the pyruvate consumption by mitochondria in MB neurons, we previously reported that LTM relies on an increased mitochondrial pyruvate flux in MB vertical lobes (a structure encompassing the axonal projections of MB neurons). This increased pyruvate metabolism occurred in the first 3 hours after the end of spaced training^23^. Since the behavioral results following Mpc1 knockdown also suggest a role for pyruvate metabolism in MTM, we performed similar imaging experiments after single-cycle training. For this, we expressed the pyruvate FRET sensor Pyronic^30^ in MB neurons (Fig. 1b) and measured pyruvate flux in the MB neurons of the vertical lobes within 1.5 hours after training. Pyruvate accumulation was measured following the application of sodium azide, a blocker of mitochondrial complex IV, as in our previously characterized method^23^. Here, we observed a faster accumulation of pyruvate in single-cycle trained flies as compared to flies that had a non-associative (unpaired) conditioning protocol, as revealed by the increased slope (Fig. 1c). This result shows that an increased mitochondrial pyruvate consumption by MB neurons occurs after single-cycle training and spaced training, in the first hours following training.

To confirm that pyruvate is actually routed to the mitochondrial TCA cycle after single-cycle training, we targeted one of the enzymes of the PDH complex, which catalyzes the conversion of pyruvate to acetyl-CoA. Knockdown of the β subunit of PDHE1 in adult MB neurons impaired MTM (Fig. 1d, Extended Data Fig. 1d) and, as previously reported using another RNAi, LTM formation (Fig. 1d, ^23^). MTM and LTM were not impaired when RNAi was not induced (Extended Data Fig. 1c-d). LT-ARM was not affected by PDHE1? knockdown (Fig. 1d). Furthermore, sensory acuity for the relevant stimuli was normal in the flies of interest (Extended Data Table 1). All together, these results show that MB neuron mitochondrial pyruvate consumption is increased for both MTM and LTM.

### Pyruvate is produced from alanine in the MB for memory formation

Next, we aimed to identify the source of neuronal pyruvate for MTM and LTM. The main routes of pyruvate production are: i) glycolysis, which allows pyruvate production from glucose through a sequence of ten enzymatic reactions; ii) lactate conversion into pyruvate, catalyzed by the lactate dehydrogenase LDH; and iii) L-alanine transamination into pyruvate, catalyzed by the alanine aminotransferase ALAT^31^ (Fig. 2a). We therefore tested each of these three routes for a role in memory formation in MB neurons. In adult MB neurons, knockdown of phosphofructokinase (PFK), an enzyme that catalyzes a rate-limiting step in glycolysis, had no effect on MTM (Fig. 2b). We previously reported that PFK knockdown in adult MB neurons had no effect on LTM, either^21^. Knockdown of LDH in adult MB neurons had no effect on MTM or LTM (Fig. 2c, Extended Data Fig. 2a). On the contrary, knocking down ALAT in adult MB neurons led to an impairment of both MTM and LTM (Fig. 2d). MTM and LTM were not impaired when RNAi was not induced (Extended Data Fig. 2b). Memory after massed training was not affected by ALAT knockdown (Fig. 2d). Sensory acuity for the relevant stimuli was normal in the flies of interest (Extended Data Table 2). We replicated these behavioral experiments using a second, non-overlapping ALAT RNAi (Extended Data Fig. 2c). Overall, these results show that MTM and LTM do not rely on neuronal glycolysis or lactate dehydrogenation, but instead rely on alanine metabolism.

**Figure 2:**
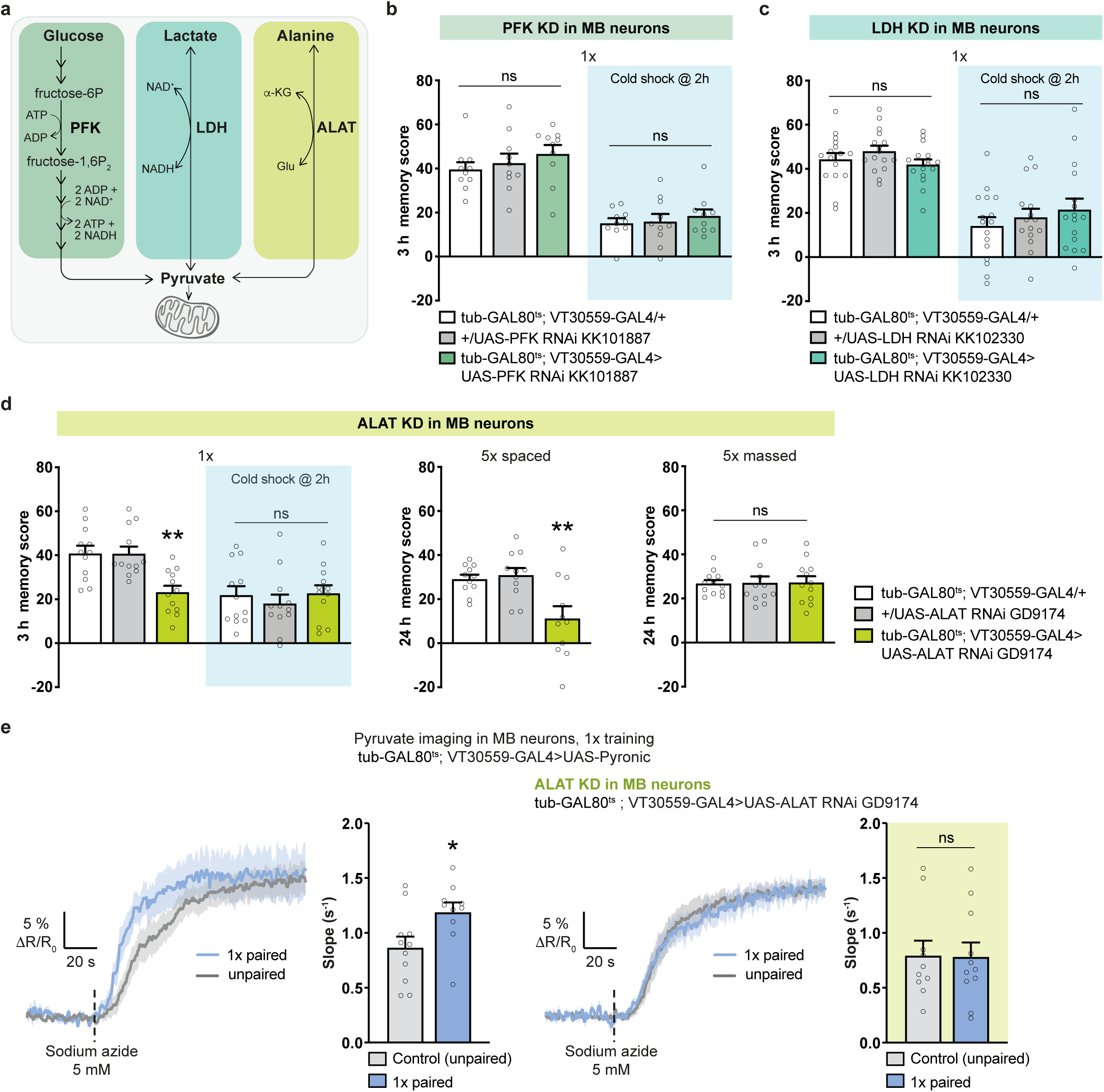
Pyruvate is produced from alanine for MTM and LTM. a. Scheme of the three main pyruvate production routes. Pyruvate can be synthesized: i) from glucose through glycolysis, including one of the rate-limiting steps catalyzed by phosphofructokinase (PFK); ii) upon lactate oxidation catalyzed by lactate dehydrogenase (LDH); and iii) upon alanine transamination catalyzed by alanine aminotransferase (ALAT). b. PFK knockdown (KD) in adult MB neurons did not affect memory after single-cycle training (n = 10, F_2,27_ = 0.85, P = 0.44) or single-cycle training followed by cold shock (n = 10, F_2,27_ = 0.35, P = 0.71). c. LDH knockdown in adult MB neurons did not affect memory after single-cycle training (n = 15-16, F_2,45_ = 1.52, P = 0.23) or single-cycle training followed by cold shock (n = 16, F_2,44_ = 0.76, P = 0.47). d. ALAT knockdown in adult MB neurons impaired memory after single-cycle training (n = 12, F_2,33_ = 10.2, P < 0.001) and spaced training (n = 11, F_2,30_ = 7.95, P = 0.002), but did not affect memory after single-cycle training followed by cold shock (n = 12, F_2,33_ = 0.41, P = 0.66) or massed training (n = 12, F_2,33_ = 0.012, P = 0.99). e. Single-cycle training elicited a faster pyruvate accumulation in MB neuron axons following sodium azide application (5 mM) as compared to non-associative unpaired training (n = 10, t_18_ = 2.27, P = 0.036). ALAT knockdown in adult MB neurons impaired the single-cycle induced increase in pyruvate accumulation in MB neuron axons following sodium azide application (n = 10-11, t_19_ = 0.064, P = 0.95). All data are presented as mean ± SEM. Asterisks illustrate the significance level of the t-test, or of the least significant pairwise comparison following an ANOVA, with the following nomenclature: *p < 0.05; **p < 0.01; ns: not significant, p > 0.05.

To confirm that alanine is indeed the source of pyruvate for MTM, we performed *in vivo* pyruvate imaging in MB vertical lobes while knocking down ALAT in adult MB neurons. Single-cycle training failed to elicit an increased pyruvate accumulation when ALAT was knocked down in adult MB neurons (Fig. 2e). Collectively, these results show that alanine is the relevant source of pyruvate in MB neurons for memory formation.

### Cortex glia transfers glycolysis-derived alanine to MB neurons for memory formation

Previously, it was shown in Drosophila that alanine can be produced as a byproduct of glial glycolysis and released by glial cells^15^. We therefore hypothesized that the alanine required for neuronal pyruvate production and consumption for MTM and LTM originates from neighboring glial cells. The transamination reaction catalyzed by ALAT is reversible, so that alanine can be produced from glycolysis-derived pyruvate. ALAT knockdown in all glial cells using repo-GAL4 impaired MTM (Fig. 3a). MTM was not impaired when RNAi was not induced (Extended Data Fig. 3a), suggesting that alanine is produced in glia for MTM. Next, we examined which specific glial cell types require ALAT. The Drosophila brain contains several types of glial cells, including three types that are in close contact with different neuronal compartments (Fig. 3b). The ensheathing glia delimit major brain structures, while astrocyte-like glial cells contact synapses and cortex glia enwrap neuron somata^32–34^. ALAT knockdown in either adult astrocyte-like glia (using the alrm-GAL4 driver) or ensheathing glia (using the 56F03-GAL4 driver) had no effect on MTM (Fig. 3c). However, ALAT knockdown in adult cortex glia (using the 54H02-GAL4 driver) impaired both MTM and LTM (Fig. 3d). MTM and LTM were not impaired when RNAi was not induced (Extended Data Fig. 3b). LT-ARM was not affected by ALAT knockdown (Fig. 3d). Sensory acuity for the relevant stimuli was normal in the flies of interest (Extended Data Table 3). We then replicated these behavioral experiments using a second, non-overlapping ALAT RNAi (Extended Data Fig. 3c).

**Figure 3:**
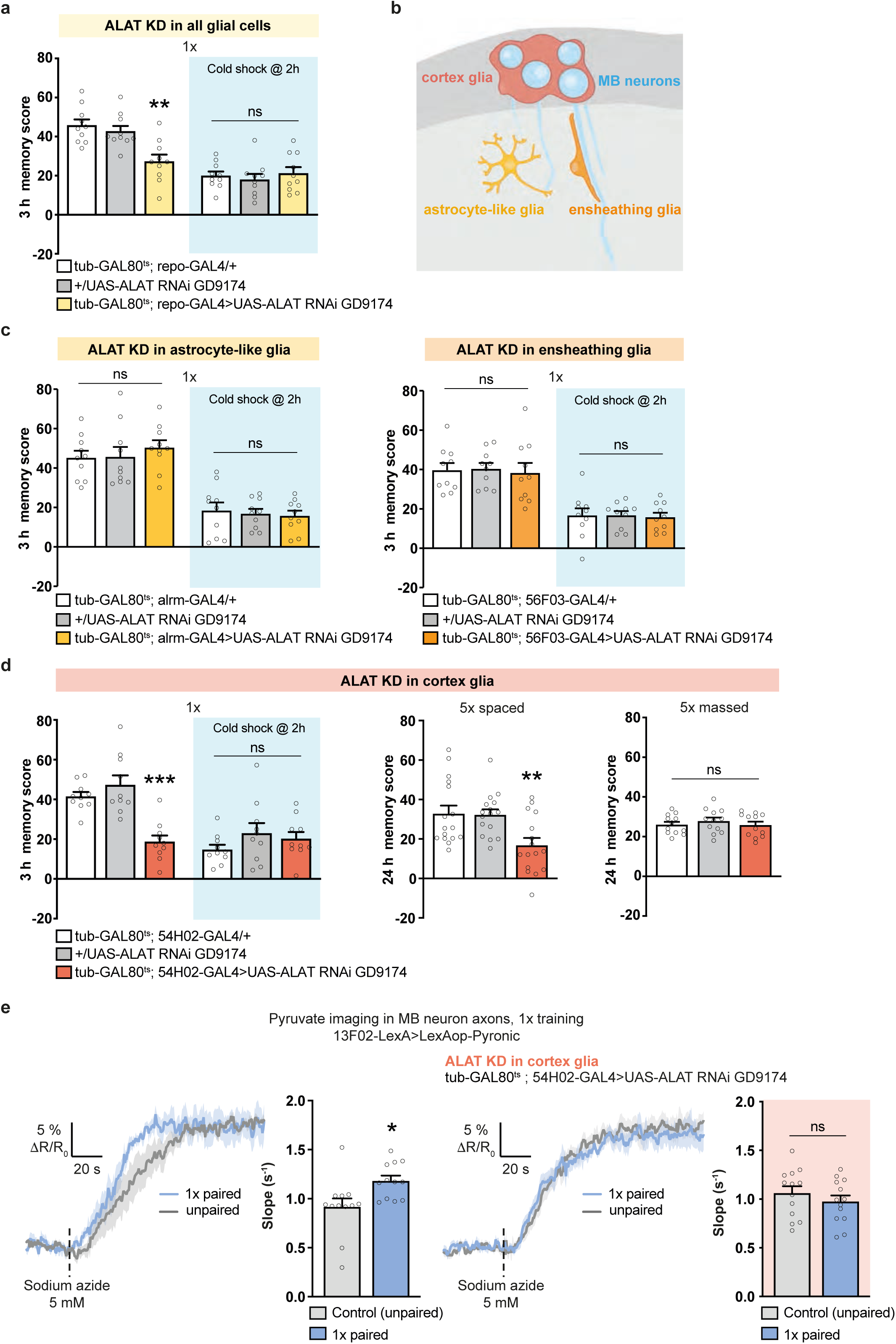
Cortex glia transfer alanine to MB neurons for MTM and LTM. a. ALAT knockdown (KD) in adult glia impaired memory after single-cycle training (n = 10, F_2,27_ = 10.56, P = 0.0004), but did not affect memory after single-cycle training followed by cold shock (n = 10, F_2,27_ = 0.38, P = 0.69). b. Subtypes and localization of Drosophila glia contacting neurons in the brain. In the cortical region (dark grey), cortex glia (red) encase the somata of neurons (blue). In the neuropile (light grey), astrocyte-like glia (yellow) extend their processes close to the synapses. The processes of ensheathing glia (orange) delimit brain anatomical structures. c. ALAT knockdown in adult astrocyte-like glia (left) or ensheathing glia (right) did not affect memory after single-cycle training (astrocyte-like glia: n = 10, F_2,27_ = 0.48, P = 0.62; ensheathing glia: n = 10, F_2,27_ = 0.07, P = 0.93) or single-cycle training followed by cold shock (astrocyte-like glia: n = 10, F_2,27_ = 0.18, P = 0.83; ensheathing glia: n = 10, F_2,27_ = 0.037, P = 0.96). d. ALAT knockdown in adult cortex glia impaired memory after single-cycle training (n = 10, F_2,27_ = 18.67, P <0.001) and spaced training (n = 16, F_2,45_ = 6.57, P = 0.0031), but did not affect memory after single-cycle training followed by cold shock (n = 10, F_2,27_ = 1.26, P = 0.30) or massed training (n = 12, F_2,33_ = 0.45, P = 0.64). e. Single-cycle training elicited a faster pyruvate accumulation in MB neuron axons following sodium azide application (5 mM) as compared to non-associative unpaired training (n = 12, t_22_ = 2.65, P = 0.015). ALAT knockdown in adult cortex glia impaired the single-cycle induced increase in pyruvate accumulation in MB neuron axons following sodium azide application (n = 13, t_24_ = 0.94, P = 0.36). All data are presented as mean ± SEM. Asterisks illustrate the significance level of the t-test, or of the least significant pairwise comparison following an ANOVA, with the following nomenclature: *p < 0.05; **p < 0.01; ***p < 0.001; ns: not significant, p > 0.05.

We hypothesized that cortex glia alanine is transferred to MB neurons as a pyruvate precursor. If so, the loss of ALAT activity solely in adult cortex glia should prevent the increased pyruvate consumption in MB neurons after training. Indeed, single-cycle training failed to elicit an increased pyruvate accumulation in the MB vertical lobes as compared to non-associative training when ALAT was knocked down in adult cortex glia (Fig. 3e). Since the cortex glia specifically surround neuronal somata, we next hypothesized that alanine is transferred to this MB neuron compartment. If so, alanine could be transaminated into pyruvate in MB neuron somata. We therefore monitored the pyruvate flux in MB neuron somata (Extended Data Fig. 3d). Sodium azide application triggered a faster pyruvate accumulation in MB neuron somata after single-cycle training as compared to non-associative training (Extended Data Fig. 3e). Furthermore, as in the axonal compartment, this effect was abolished by ALAT knockdown in adult cortex glia (Extended Data Fig. 3e). Together, our behavioral and imaging results suggest that the cortex glia produce alanine to sustain neuronal pyruvate needs during MTM and LTM formation.

Next, we sought to confirm whether alanine production is derived from glucose metabolism in cortex glia. Indeed, knockdown of the glycolytic enzyme PFK in adult cortex glia impaired both MTM and LTM (Fig. 4a). MTM and LTM were not impaired when RNAi was not induced (Extended Data Fig. 4a). LT-ARM was not affected by PFK knockdown (Fig. 4a). Sensory acuity for the relevant stimuli was normal in the flies of interest (Extended Data Table 4). These behavioral experiments were replicated using a second, non-overlapping PFK RNAi (Extended Data Fig. 4b). In addition, single-cycle training failed to elicit increased pyruvate consumption in the MB vertical lobes or somata as compared to non-associative training when PFK was knocked down in adult cortex glia (Fig. 4b, Extended Data Fig. 4c). These behavioral and imaging results therefore suggest that cortex glia glycolysis is required for alanine production as well as MTM and LTM formation. Collectively, these data demonstrate the existence of a cortex glia-to-MB neuron alanine transfer that is dependent on glial glycolysis to sustain their pyruvate need required by memory formation.

**Figure 4:**
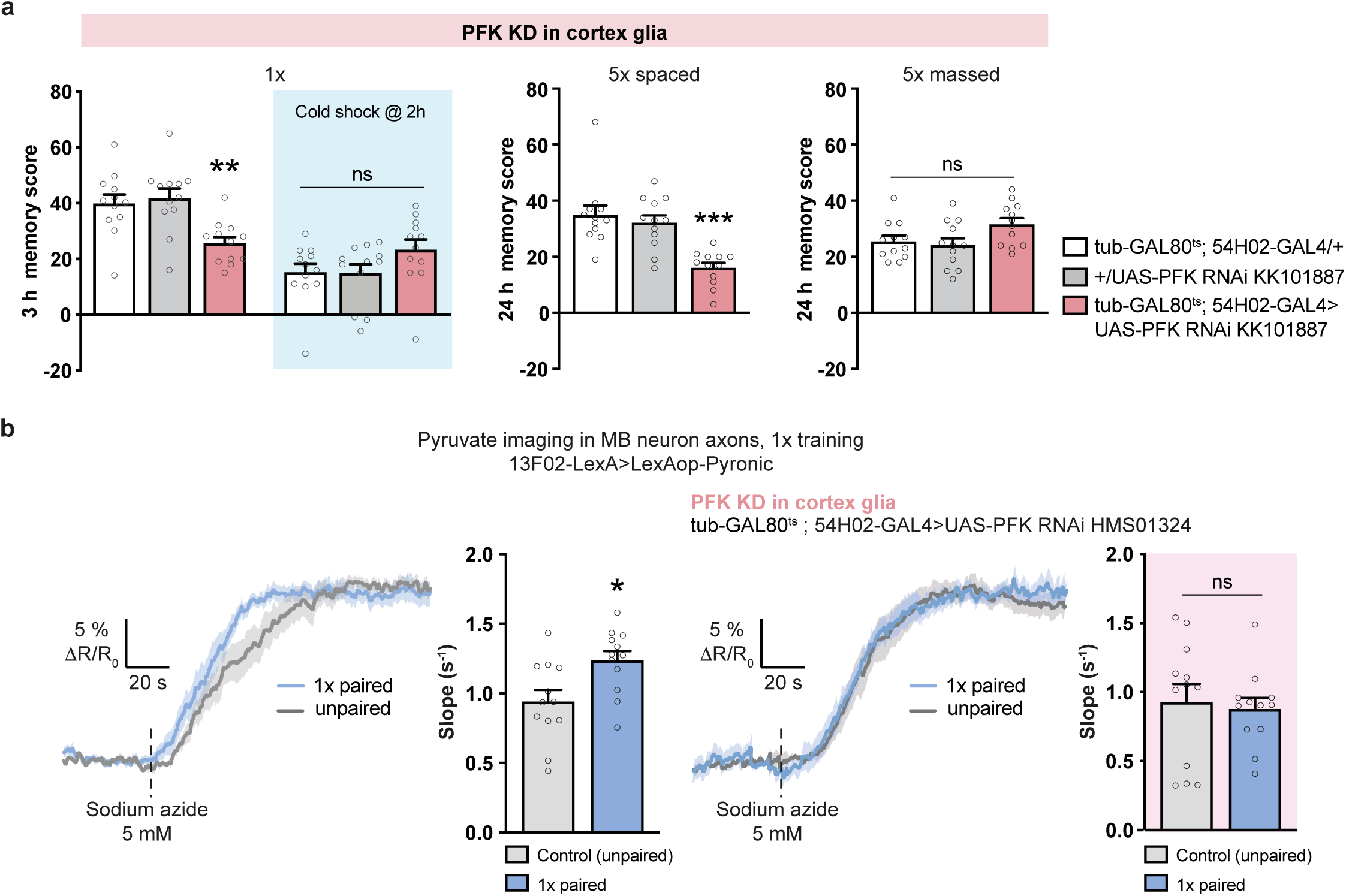
Glycolysis in cortex glia is necessary for MTM and LTM. a. PFK knockdown (KD) in adult cortex glia impaired memory after single-cycle training (n = 12, F_2,33_ = 8.26, P = 0.001) and spaced training (n = 12, F_2,33_ = 14.21, P <0.001), but did not affect memory after single-cycle training followed by cold shock (n = 12, F_2,33_ = 1.99, P = 0.15) or massed training (n = 12, F_2,33_ = 3.16, P = 0.055). b. Single-cycle training elicited a faster pyruvate accumulation in MB neuron axons following sodium azide application (5 mM) as compared to non-associative unpaired training (n = 12, t_22_ = 2.75, P = 0.012). PFK knockdown in adult cortex glia impaired the single-cycle induced increase in pyruvate accumulation in MB neuron axons following sodium azide application (n = 12, t_22_ = 0.34, P = 0.74). All data are presented as mean ± SEM. Asterisks illustrate the significance level of the t-test, or of the least significant pairwise comparison following an ANOVA, with the following nomenclature: *p < 0.05; **p < 0.01; ns: not significant, p > 0.05.

### Cortex glia take up glucose through the transporter glug to produce alanine

Our results show that glycolysis in cortex glia is required for MTM and LTM. Cortex glia have multiple potential sources of glucose. Indeed, cortex glia can: i) import trehalose as they express the enzyme trehalase^35^, which catalyzes the breakdown of trehalose into glucose; ii) directly take up glucose from the extracellular space^21,36^ to fuel glycolysis; or iii) use glucose-6-phosphate derived from glycogen breakdown^37^. Therefore, we aimed to decipher the origin of glucose in the fueling of glycolysis for MTM formation. Trehalase knockdown in adult cortex glia had no effect on MTM (Extended Data Fig. 5a). Similarly, knocking down glycogen phosphorylase (GlyP), which catalyzes the breakdown of glycogen into glucose-1-phosphate, in adult cortex glia had no effect on MTM (Extended Data Fig. 5b). These results suggest that the glucose used for glycolysis-derived alanine production might be directly imported into the cortex glia through a glucose transporter. We previously published that Drosophila Glut1, the homolog of mammalian members of the SLC2 membrane transporters Glut1 and Glut3^38^, is responsible for glucose export from cortex glia upon LTM formation, rather than being responsible for glucose import^21^. According to single-cell transcriptomic data, several other members of the SLC2 family of glucose transporters are highly expressed in glial cells^35^. Specifically, among the SLC2 family, two genes encoding predicted hexose sugar transporters are enriched in cortex glia as compared to other transporters: CG31100, that we named ‘glug’ (for <u>gl</u>ucose <u>u</u>ptake by <u>g</u>lia) and nebu (Extended Data Fig. 5c). They represent therefore good candidates for mediating glucose uptake in cortex glia to sustain MTM and LTM. Knockdown of nebu in adult cortex glia had no effect on MTM (Fig. 5a). Interestingly, knockdown of glug in adult cortex glia led to MTM and LTM defects, whereas memory after massed training remained intact (Fig. 5b). MTM and LTM were not impaired when RNAi was not induced (Extended Data Fig. 5d). Sensory acuity for the relevant stimuli was normal in the flies of interest (Extended Data Table 5). Next, to test whether glug mediates the glucose uptake necessary for glycolysis-derived alanine production aimed at fueling MB neurons, we performed *in vivo* pyruvate imaging in MB neurons while knocking down glug in cortex glia. As seen for PFK and ALAT, we found that glug knockdown in adult cortex glia abolished the increase in pyruvate consumption in MB neuron vertical lobes induced by single-cycle training (Fig. 5c) as well as in MB neuron somata (Extended Data Fig. 5e). Conversely, when nebu was knocked down in adult cortex glia, we were still able to observe an increase in pyruvate flux after single-cycle training (Fig. 5c, Extended Data Fig. 5e). Together, these data show that sugar import into cortex glia *via* glug is necessary for glial glycolysis supporting neuronal pyruvate metabolism.

**Figure 5:**
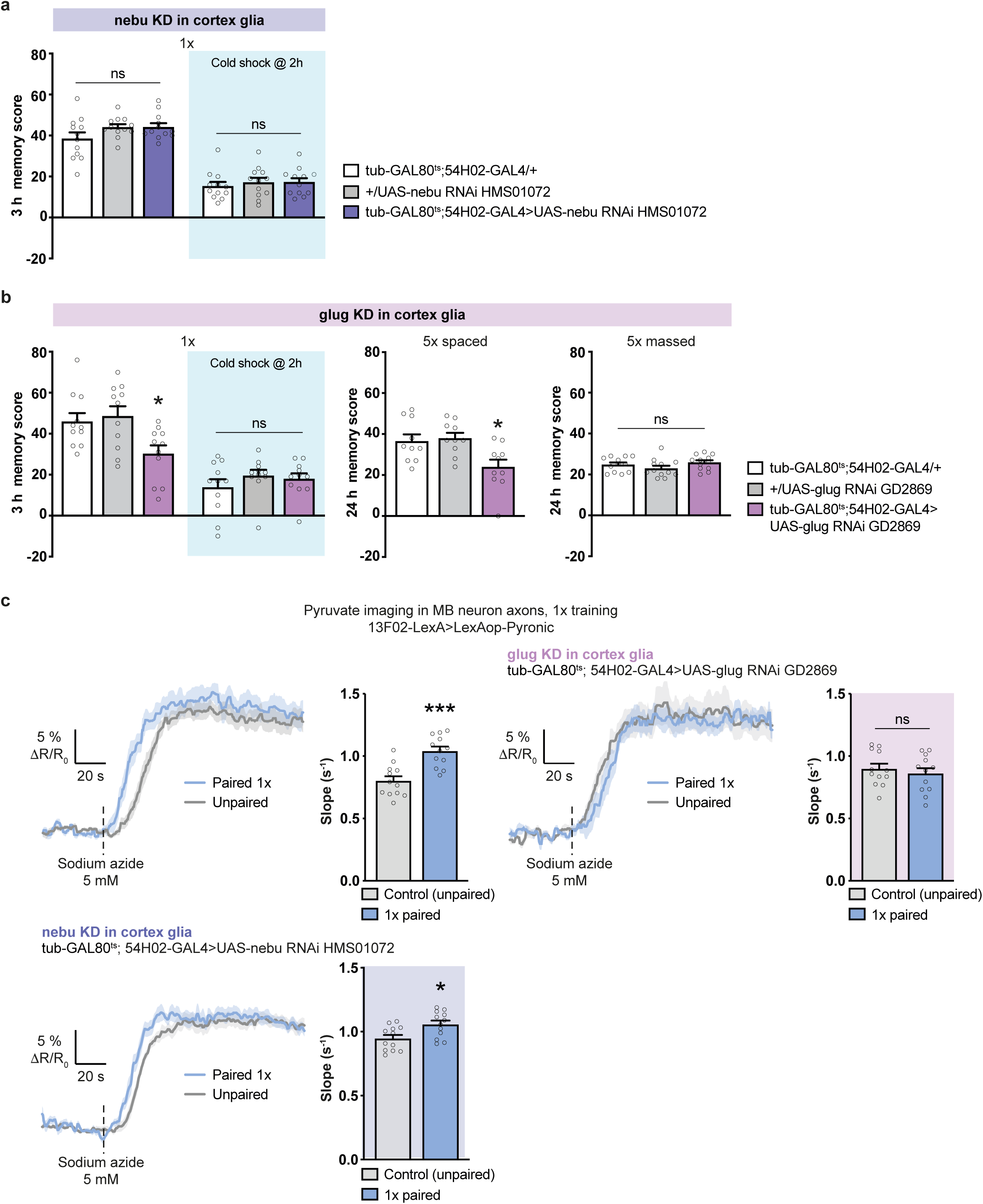
The sugar transporter glug is required in cortex glia for glycolysis-derived alanine synthesis specifically. a. Nebu knockdown (KD) in adult cortex glia did not affect memory after single-cycle training (n = 12, F_2,33_ = 2.21, P = 0.13) or single-cycle training followed by cold shock (n = 12, F_2,33_ = 0.31, P = 0.73) b. Glug knockdown in adult cortex glia impaired memory after single-cycle training (n = 11, F_2,30_ = 5.36, P = 0.01) and spaced training (n = 10, F_2,27_ = 5.92, P = 0.007), but did not affect memory after single-cycle training followed by cold shock (n = 11, F_2,30_ = 0.84, P = 0.44) or massed training (n = 12, F_2,33_ = 1.73, P = 0.19). c. Single-cycle training elicited a faster pyruvate accumulation in MB neuron axons following sodium azide application (5 mM) as compared to non-associative unpaired training (n = 12, t_22_ = 4.62, P<0.001). Glug knockdown in adult cortex glia impaired the single-cycle induced increase in pyruvate accumulation in MB neuron axons following sodium azide application (n = 12, t_22_ = 0.65, P = 0.52). When nebu was knocked down in adult cortex glia, single-cycle training continued to elicit an increase in pyruvate accumulation in MB neuron axons following sodium azide application (n = 12, t_22_ = 2.73, P = 0.012). All data are presented as mean ± SEM. Asterisks illustrate the significance level of the t-test, or of the least significant pairwise comparison following an ANOVA, with the following nomenclature: *p < 0.05; ***p < 0.001; ns: not significant, p > 0.05.

### Cortex glia take up glucose through nebu and transfer it to MB neurons

In addition to providing alanine to fuel neuronal mitochondria for MTM and LTM, cortex glia glucose is involved in a metabolite exchange with neurons that is specific to LTM formation^21^. Indeed, we previously demonstrated that cortex glia activation through the nAchRα7 nicotinic receptor triggers an insulin-dependent increase in glucose concentration, allowing the transfer of glucose to MB neurons to sustain neuronal PPP, specifically for LTM formation^21^. As the glucose transporter mediating glucose import in cortex glia for LTM was not identified at that time, we wondered if the transporter identified here, glug, could also be involved in this process. To test this hypothesis, we performed *in vivo* glucose imaging experiments in cortex glia in response to nicotine stimulation, using a FRET glucose sensor (Fig. 6a) and our previously well-characterized method^21^. Nicotine application resulted in the expected rise of glucose concentration in cortex glia, whereas glug knockdown in adult cortex glia did not affect the nicotine-induced glucose increase (Fig. 6b). To confirm that glug does not mediate acetylcholine-dependent glucose import into cortex glia, we checked that it was not involved in the LTM-specific cortex glia-to-MB neuron glucose shuttle. For this, we performed glucose imaging in MB neuron somata (Fig. 6c) and monitored glucose consumption after spaced training, as previously described^21^. We thus confirmed that spaced training triggers increased glucose consumption by MB neuron somata in normal flies. A similar increase in glucose consumption by MB neuron somata was observed when glug was knocked down in cortex glia (Fig. 6d). Together, these data show that glug does not mediate glucose uptake for the LTM-specific cortex glia-MB neuron glucose shuttle.

**Figure 6:**
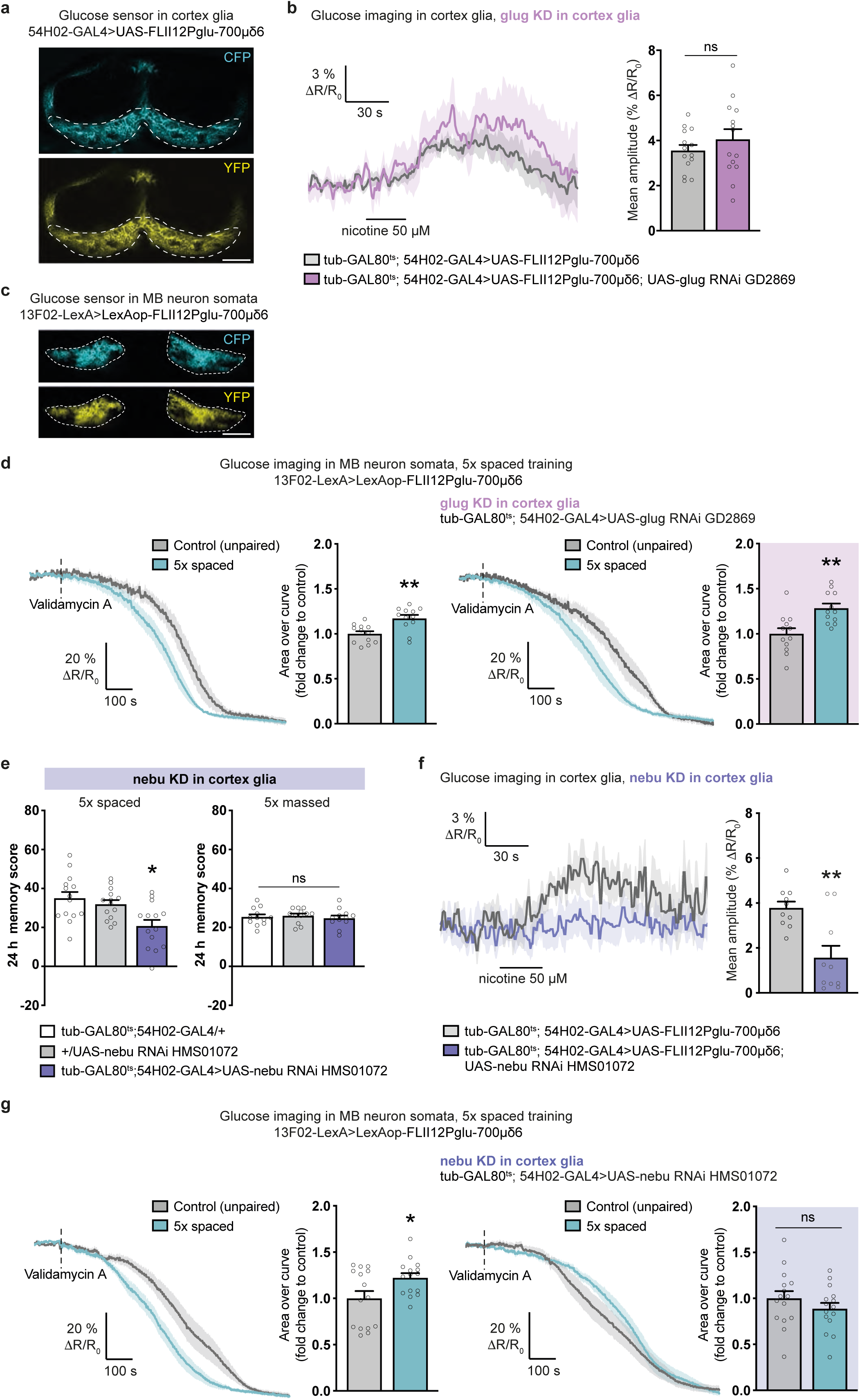
A parallel glucose import into cortex glia through nebu allows direct glucose transfer to MB neurons. a. The FRET glucose sensor FLII12Pglu-700μδ6^58^ was expressed in adult cortex glia. The two images show the CFP and YFP channels. The dashed lines delimit the cortex glia region where the glucose FRET signal was quantified. Scale bar: 50 µm. b. Nicotine stimulation (50 µM, 30 s) increased glucose concentration in cortex glia. Knockdown (KD) of glug in adult cortex glia had no effect on the nicotine-induced glucose elevation (n = 14, t_26_ = 0.99, P = 0.33). c. The FRET glucose sensor FLII12Pglu-700μδ6 was expressed in adult MB neurons. The two images show the CFP and YFP channels. The dashed lines delimit MB neuron somata where the glucose FRET signal was quantified. Scale bar: 30 µm. d. The glucose concentration in MB neuron somata following the application of validamycin A (4 mM, dashed line) decreased faster in flies after spaced training as compared to flies conditioned with a non-associative spaced unpaired training protocol (n = 12, t_22_ = 3.56, P = 0.002). When glug was knocked down in adult cortex glia, spaced training continued to elicit a faster decrease of glucose concentration in MB neurons as compared to unpaired controls (n = 12, t_22_ = 3.52, P = 0.002). e. Nebu knockdown in adult cortex glia impaired memory after spaced training (n = 14, F_2,39_ = 6.92, P = 0.003), but did not affect memory after massed training (n = 12, F_2,33_ = 0.25, P = 0.78). f. Nicotine stimulation (50 µM, 30 s) increased glucose concentration in cortex glia. Knockdown of nebu in adult cortex glia impaired the nicotine-induced glucose elevation (n = 10, t_18_ = 3.70, P = 0.002). g. The glucose concentration in MB neuron somata following the application of validamycin A (4 mM, dashed line) decreased faster in flies after spaced training as compared to flies conditioned with a non-associative spaced unpaired training protocol (n = 15, t_28_ = 2.38, P = 0.025). Spaced training failed to elicit a faster decrease of glucose in MB neurons upon nebu knockdown in adult cortex glia (n = 15, t_28_ = 1.10, P = 0.28). All data are presented as mean ± SEM. Asterisks illustrate the significance level of the t-test, or of the least significant pairwise comparison following an ANOVA, with the following nomenclature: *p < 0.05; **p < 0.01; ns: not significant, p > 0.05.

We therefore hypothesized that another glucose transporter might mediate glucose uptake in cortex glia specifically for LTM. Knockdown of nebu in adult cortex glia impaired LTM (Fig. 6e). LTM was not impaired when RNAi was not induced (Extended Data Fig. 6a). Sensory acuity for the relevant stimuli was normal in the flies of interest (Extended Data Table 6). These results were confirmed with a second, non-overlapping RNAi (Extended Data Fig. 6b). Since nebu is specifically required for LTM, we tested whether it mediates the nicotine-induced glucose uptake in cortex glia. Indeed, knocking down nebu in adult cortex glia dampened the nicotine-induced glucose increase in cortex glia (Fig. 6f). Furthermore, spaced training failed to induce any glucose consumption increase in MB neurons when nebu was knocked down in cortex glia (Fig. 6g). Collectively, these results reveal that glial glycolysis for MTM and LTM and LTM-specific glucose import are supported by two different transporters in cortex glia, i.e. glug and nebu.

## Discussion

In this study, we investigated how glial-neuron metabolic coupling meets the energy demands of memory formation in Drosophila. We found that glucose drawn from the extracellular space by cortex glia is used to produce glycolysis-derived alanine. Alanine is in turn used by MB neurons as an energy substrate, which fuels the increased pyruvate mitochondrial uptake occurring after both single-cycle and spaced training and sustains the formation of MTM and LTM. Combined, these processes constitute a cortex glia-MB neuron alanine transfer at the root of memory formation (Fig. 7). We revealed that on top of glucose supporting the alanine shuttle, the cortex glia mobilize, *via* the nebu transporter, an additional flow of glucose specifically for LTM, fueling the PPP in MB neurons^21^ (Fig. 7). The model we describe raises the following questions: Why is alanine used as an alternative energy substrate for memory instead of lactate, which has been described as necessary for memory formation in rats^8^? And is alanine used as an energy substrate in other contexts?

**Figure 7:**
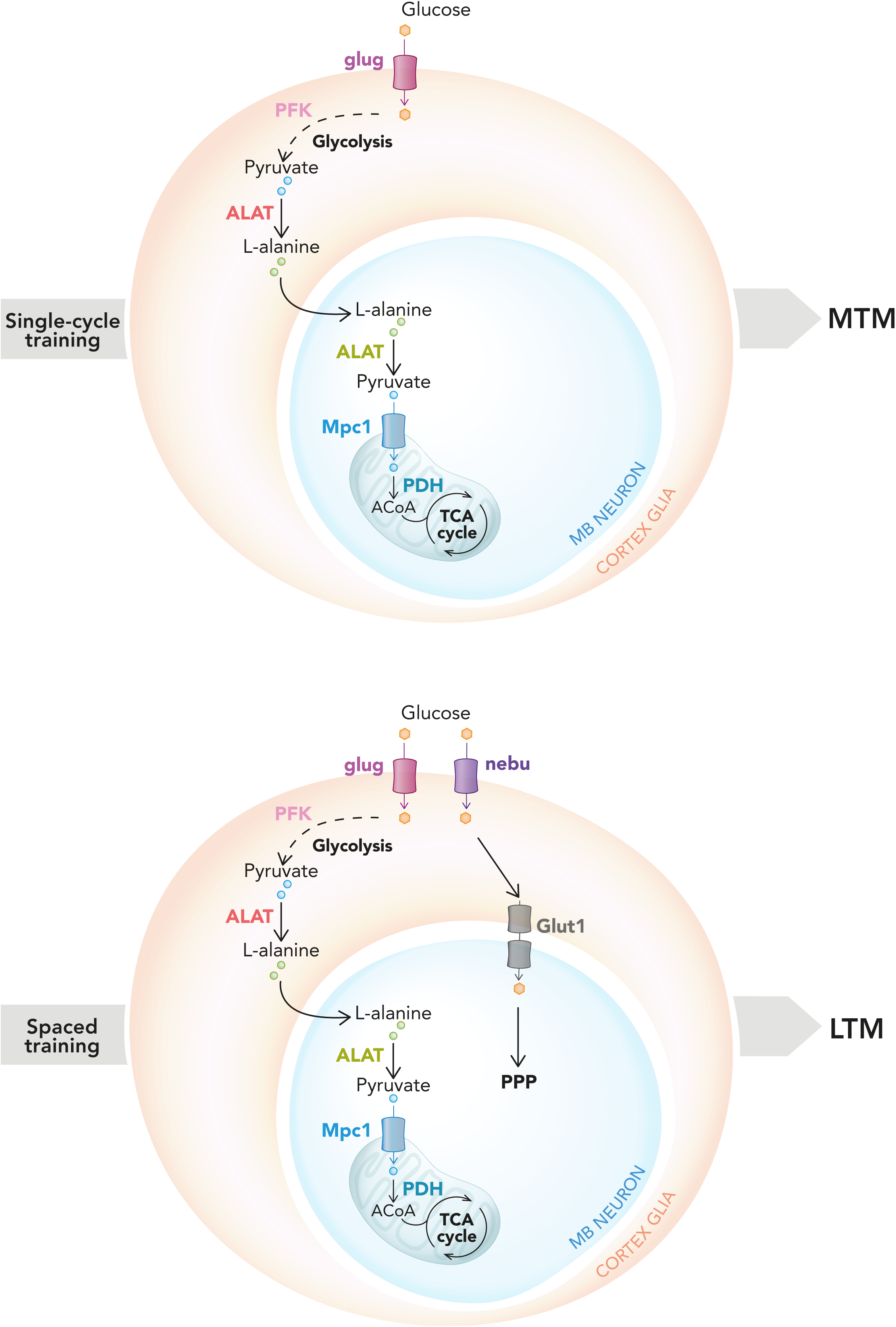
Model of cortex glia – MB neuron alanine shuttling for MTM and LTM. Upon MTM and LTM formation, hexose sugar, most likely glucose, taken up by cortex glia through glug is routed to glycolysis, including one reaction catalyzed by phosphofructokinase (PFK) to form pyruvate. Pyruvate is then transaminated to alanine by alanine aminotransferase (ALAT). Alanine in MB neurons is transaminated back to pyruvate by ALAT. Pyruvate transported into mitochondria through mitochondrial pyruvate carrier 1 (Mpc1) is transformed into acetyl-CoA (ACoA) by pyruvate dehydrogenase (PDH), where it is likely integrated into the tricarboxylic acid cycle (TCA) cycle to produce energy upon oxidative phosphorylation. Upon LTM formation, the alanine shuttle is supplemented by the glucose shuttle. Glucose entering cortex glia through nebu is directly shuttled to MB neurons through Glut1, upon a nicotinic-insulinergic activation cascade in cortex glia (see ^21^). Neuronal glucose is then routed to the pentose phosphate pathway (PPP).

Several pieces of evidence point to the potential use of alanine as a neuronal energy substrate in other species. In PDH-deficient mice, which have disrupted mitochondrial metabolism, NMR spectroscopy analysis has revealed the accumulation of alanine in the brain, suggesting that it is a substrate that normally fuels mitochondrial metabolism^39^. In support of this study, it was confirmed *ex vivo* that alanine can be produced from glucose in the murine hippocampus^40^, although the cellular origin of alanine was not identified. ALAT is expressed by both neurons and astrocytes in the mouse and human brain^41,42^. Measurements of metabolic activity of cultured cortical neurons has revealed substantial alanine uptake from neurons^17^. Moreover, cultured cortical neurons produce CO_2_ from alanine, indicating that they can metabolize it for energy purposes. In parallel, cultured astrocytes release alanine produced from glucose^17,43^. These features are consistent with the existence of an alanine transfer from glial cells to neurons, at least in basal conditions. Our study takes this further by showing that this alanine shuttling is physiologically relevant and specifically activated *in vivo* upon memory formation. In support of our data, previous NMR studies have demonstrated that glucose-derived alanine is enriched in activated areas of the rat brain upon sensory stimulation^44,45^. An activity-dependent glia-to-neuron alanine shuttle was further described in studies using *ex vivo* honeybee retina preparations, in which stimulation of photoreceptors triggered an increase in alanine use for energy production^46^. In the same model, the alanine source is the neighboring glial cells^16^. A similar system exists in the mammalian retina in which specialized astrocytes, namely Müller cells, transaminate pyruvate into alanine in response to glutamate stimulation. Collectively, these studies support the existence of a glia-neuron alanine shuttle across several species, similar to the cortex glia-MB neuron shuttle that we have described for memory formation in Drosophila. Notably, neither these studies nor our present work have identified the amino acid transporter that mediates the glia-to-neuron alanine transfer. In humans, several ALAT mutations have been associated with intellectual disability, revealing the importance of alanine metabolism for cerebral functions^47–49^. Lastly, increased alanine metabolism has been correlated with the development of the nervous system. Indeed, ALAT expression is increased at the peaks of synaptogenesis in the developing mouse hippocampus^49^. Additionally, the ALAT knockout mouse model exhibits decreased TCA cycle activity associated with a decreased brain size and reduced synapse formation^49^. This supports the participation of the glia-neuron alanine shuttle in other energy-costly cerebral processes than memory.

By monitoring neuronal pyruvate consumption *in vivo*, we previously demonstrated that pyruvate metabolism in neurons is increased following LTM formation^23^. This raised the question of the nature of the fuel enabling pyruvate metabolism in MB neurons. In a more recent study, we ruled out neuronal glycolysis as the source of MB neuron pyruvate for LTM^21^. Here, we demonstrated that pyruvate uptake in MB neuron mitochondria was necessary for both LTM and MTM. The upregulation of pyruvate metabolism in MB neurons depends on ALAT activity, suggesting that alanine is the substrate for pyruvate production. Because the knockdown of every gene we described to be involved in the cortex glia-MB neuron alanine shuttle pathway systematically impaired both MTM and LTM, it is likely that glial alanine also fuels the increased pyruvate metabolism of MB neurons occurring during LTM formation. In contrast, studies conducted in rats have shown that LTM depends on lactate import from astrocytes^8^. This brings into question what advantage producing alanine instead of lactate could offer. It should be noted that the ANLS model was initially proposed to account for astrocyte neuron coupling at glutamatergic synapses. Glycolysis and lactate production in astrocytes are directly coupled, as lactate dehydrogenase enzyme (LDH) regenerates NAD^+^ from the NADH produced during glycolysis. In parallel, astrocytes can oxidize glutamate to produce NADH as well as TCA cycle intermediates to sustain their own energy needs^50,51^. Indeed, glutamate is a significant energy substrate in astrocytes as it contributes to more than 20% of astrocytic oxidative metabolism^52,53^. In non-glutamatergic systems, it is likely that glial cells need alternative sources to sustain their energy metabolism. In the absence of LDH activity, glycolytic NADH can be spared for glial metabolic needs. Furthermore, the transamination reaction of pyruvate into alanine catalyzed by ALAT does not require the conversion of NADH to NAD^+^. Therefore, producing alanine instead of lactate in glial cells would allow saving glycolysis-derived NADH for glial ATP production. This would make the alanine shuttle an efficient mechanism to sustain the energy needs of glial cells and neighboring non-glutamatergic neurons. In support of such a neuron subtype-specific utilization of alanine, studies conducted *in vitro* show that GABAergic neurons display a higher net alanine uptake rate than glutamatergic neurons^43^. Here, we describe a model in which alanine is the end-product of glial glycolysis, fueling cholinergic MB neurons. We therefore hypothesize that the alanine shuttle is a relevant mechanism sustaining the energy needs of cholinergic neurons in general.

We demonstrated that alanine is produced by cortex glial cells upon glycolysis. Contrary to the similar lactate shuttle occurring for memory in rats^8^, we showed that glycogen breakdown is not the source of glucose for glia in Drosophila. Instead, we identified glug, a member of the SLC2 hexose sugar transporters which are highly expressed in cortex glia^35^, as the hexose transporter in cortex glia for MTM and LTM. Glug also mediates the increased pyruvate consumption in MB neurons accompanying MTM formation, consistent with a role in importing sugar for use in glycolysis-derived alanine production.

We previously revealed the existence of a nicotinic acetylcholine receptor-dependent glucose shuttle from cortex glia to MB neurons that fuels the neuronal PPP, crucial for LTM formation^21^. As glug neither mediated this nicotine-induced glucose increase in cortex glia nor the subsequent increased glucose consumption by MB neurons used for LTM, we searched for a different glucose transporter in cortex glia that would be specifically required for LTM. We thus identified nebu, another member of the SLC2 family that is also highly expressed in cortex glia according to single-cell transcriptomic profiling in the fly brain^35^, as necessary for LTM specifically. In addition, nebu was necessary for both nicotine-induced glucose uptake in cortex glia and the increased glucose consumption in MB neurons following LTM conditioning. Interestingly, we previously reported that the glucose shuttle from cortex glia to MB neurons for LTM was dependent on insulin signaling^21^. Indeed, the glucose concentration increase in cortex glia depends on insulin peptide Ilp4, which acts autocrinally on its receptor InR in cortex glia. Insulin-dependent glucose uptake has been reported in mammalian peripheral organs and the brain, where it acts through the insulin-sensitive glucose transporter Glut4^54,55^. Similar to what occurs in mammalian adipose tissue in response to insulin, InR signaling in Drosophila fat cells allows glucose uptake and storage^56,57^. However, to date, no equivalent glucose transporter responding to insulin has been identified in Drosophila^56^. Here, we demonstrate that the glucose transporter nebu and the insulin receptor belong to the same pathway and act together to increase glucose concentration in cortex glia, suggesting that nebu functions as an insulin-sensitive glucose transporter in Drosophila.

Overall, this work, along with our recent demonstration that glia provide glucose to neurons for the PPP, provides a global picture of glucose metabolism underlying memory formation in Drosophila, highlighting the existence of two parallel fates for glucose orchestrated by cortex glia. Whereas the glycolytic production of alanine sustains both MTM and LTM, direct glucose transfer to neurons for the PPP is a specific feature of LTM formation. It is intriguing that these two parallel fates involve distinct glucose transporters in an apparently independent fashion. Indeed, glug knockdown prevented glucose transfer after spaced training whereas nebu knockdown did not alter alanine production, suggesting that the activities of the two transporters do not simply add up. Instead, more sophisticated mechanisms are probably involved, which may involve separate cellular pools of glucose or the coupling of enzyme clusters with specific transporters, warranting further investigation.

## Methods

### Experimental model

Flies (*Drosophila melanogaster*) were raised on standard medium at 18 or 23°C (depending on the experiments; see respective details below) and 60% humidity in a 12-h light/dark cycle. The study was performed on 1-4-day-old adult flies. For behavior experiments, both male and female flies were used. For imaging experiments, female flies were used because of their larger size. All flies obtained from libraries or received after the injection of transgenes (except flies from the TRiP RNAi collection) were outcrossed for 5 generations to a reference strain carrying the w^1118^ mutation in an otherwise Canton Special (Canton S) genetic background. Because TRiP RNAi transgenes are labelled by a y^+^ marker, these lines were outcrossed to a y^1^w^67c23^ strain in an otherwise Canton S background. All strains used in this study are described in Extended Data Table 7.

### Behavior Experiments

For behavior experiments, flies were raised on standard medium at 18°C and 60% humidity in a 12-h light/dark cycle. We used the TARGET system^29^ to inducibly express RNAi constructs exclusively in adult flies and not during development. To achieve the induction of RNAi expression, adult flies were kept at 30.5°C for 3 days before conditioning. Otherwise, experimental flies (0-3 days old) were transferred to fresh bottles containing standard medium 24 h before conditioning.

The behavior experiments, including the sample sizes, were conducted similarly to other studies from our laboratory^21,23,59^. Groups of 20-50 flies were subjected to one of the following olfactory conditioning protocols: a single cycle (1x training), five consecutive associative training cycles (5x massed training), or five associative cycles spaced by 15-min inter-trial intervals (5x spaced training). Non-associative control protocols (unpaired protocols) were also employed for imaging experiments. Conditioning was performed using previously described barrel-type machines that allow the parallel training of up to 6 groups. Throughout the conditioning protocol, each barrel was plugged into a constant air flow at 2 L.min^-1^. For a single cycle of associative training, flies were first exposed to an odorant (the CS^+^) for 1 min while 12 pulses of 5-s long 60 V electric shocks were delivered; flies were then exposed 45 s later to a second odorant without shocks (the CS-) for 1 min. The odorants 3-octanol and 4-methylcyclohexanol, diluted in paraffin oil to a final concentration of 2.79 ·10^−1^ g.L^-1^, were alternately used as conditioned stimuli. During unpaired conditionings, the odor and shock stimuli were delivered separately in time, with shocks occurring 3 min before the first odorant.

Flies were kept on standard medium between conditioning and the memory test, either at 25°C for flies tested 3 h after training or at 18°C for flies tested 24 h after training. To test for anesthesia-resistant memory after 1x training, flies were subjected to cold treatment exposure (4°C for 2 min) 1 h before testing. The memory test was performed in a T-maze apparatus, typically 3 h after single-cycle training or 24 h after massed or spaced training. Each arm of the T-maze was connected to a bottle containing 3-octanol and 4-methylcyclohexanol, diluted in paraffin oil to a final concentration identical to the one used for conditioning. Flies were given 1 min to choose between either arm of the T-maze. A performance score was calculated as the number of flies avoiding the conditioned odor minus the number of flies preferring the conditioned odor, divided by the total number of flies. A single performance index value is the average of two scores obtained from two groups of genotypically identical flies conditioned in two reciprocal experiments, using either odorant (3-octanol or 4-methylcyclohexanol) as the CS^+^. The indicated ‘n’ is the number of independent performance index values for each genotype.

The shock response tests were performed at 25°C by placing flies in two connected compartments; electric shocks were provided in only one of the compartments. Flies were given 1 min to move freely in these compartments, after which they were trapped, collected, and counted. The compartment where the electric shocks were delivered was alternated between two consecutive groups. Shock avoidance was calculated as for the memory test.

Because the delivery of electric shocks can modify olfactory acuity, our olfactory avoidance tests were performed on flies that had first been presented another odor paired with electric shocks. Innate odor avoidance was measured in a T-maze similar to those used for memory tests, in which one arm of the T-maze was connected to a bottle with odor diluted in paraffin oil and the other arm was connected to a bottle with paraffin oil only. Naive flies were given the choice between the two arms during 1 min. The odor-interlaced side was alternated for successively tested groups. Odor concentrations used in this assay were the same as for the memory assays. At these concentrations, both odorants are innately repulsive.

### *In vivo* pyruvate imaging

Pyruvate imaging experiments were performed on flies expressing the pyruvate sensor in MB neurons *via* the VT30559-GAL4 driver or 13F02-LexA driver, in combination with either UAS-Pyronic^23^ or LexAop-Pyronic (this study). RNAis were expressed in MB neurons using the inducible tub-GAL80ts; VT30559-GAL4 driver, or in cortex glia using the inducible tub-GAL80ts; 54H02-GAL4 driver. For imaging experiments, flies were raised at 23°C to increase the expression level of genetically encoded sensors. To achieve the induction of RNAi expression, adult flies were kept at 30.5°C for 3 days before conditioning.

As in all previous imaging work from our laboratory, all *in vivo* imaging was performed on female flies, which are preferred since their larger size facilitates surgery. Data were collected indiscriminately from 30 min to 1.5 h after 1x training. A single fly was picked and prepared for imaging as previously described^60^. Briefly, the head capsule was opened and the brain was exposed by gently removing the superior tracheae. The head capsule was bathed in artificial hemolymph solution for the duration of the preparation. The composition of this solution was: NaCl 130 mM (Sigma cat. # S9625), KCl 5 mM (Sigma cat. # P3911), MgCl_2_ 2 mM (Sigma cat. # M9272), CaCl_2_ 2 mM (Sigma cat. # C3881), D-trehalose 5 mM (Sigma T cat. # 9531), sucrose 30 mM (Sigma cat. # S9378), and HEPES hemisodium salt 5 mM (Sigma cat. # H7637). At the end of surgery, any remaining solution was absorbed and a fresh 90-μL droplet of this solution was applied on top of the brain. Two-photon imaging was performed using a Leica TCS-SP5 upright microscope equipped with a 25x, 0.95 NA water-immersion objective. Two-photon excitation was achieved using a Mai Tai DeepSee laser tuned to 825 nm. The frame rate was 1 or 2 images per second.

Measurements of pyruvate consumption were performed according to a previously well-characterized protocol^23^. After 1 min of baseline acquisition, 10 µl of a 50 mM sodium azide solution (Sigma cat. #71289; prepared in the same artificial hemolymph solution) were injected into the 90-µl droplet bathing the fly’s brain, bringing sodium azide to a final concentration of 5 mM.

For the analysis of all pyruvate imaging experiments, regions of interest (ROI) were delimited by hand around each visible MB vertical lobes or MB neuron somata, and the average intensity of the mTFP and Venus channels over each ROI was calculated over time after background subtraction. The Pyronic sensor was designed so that FRET from mTFP to Venus decreases when pyruvate concentration increases. To obtain a signal that positively correlates with pyruvate concentration, the inverse FRET ratio was computed as mTFP intensity divided by Venus intensity. This ratio was normalized by a baseline value calculated over the 30 s preceding drug injection. The slope was calculated between 10 and 70% of the plateau, while the rise time was calculated from t_0_ to the time reaching 70% of the plateau. The indicated ‘n’ is the number of animals that were assayed in each condition.

### *In vivo* glucose imaging

Glucose imaging experiments were performed on flies expressing the glucose sensor in cortex glia using the 54H02-GAL4 driver or in MB neurons *via* the 13F02-LexA driver, in combination with either UAS-FLII12Pglu-700μδ6^23^ or LexAop-FLII12Pglu-700μδ6^23^. RNAis were expressed in cortex glia using the inducible tub-GAL80ts; 54H02-GAL4 driver. Fly preparation was performed as described for *in vivo* pyruvate imaging. Two-photon excitation was achieved using a Mai Tai DeepSee laser tuned to 820 nm. The frame rate was 1 image per second.

Nicotine stimulation experiments were performed as previously described^21^. On each experimental day, nicotine was freshly diluted from a commercial liquid (Sigma N3876) into the saline used for imaging. A perfusion setup at a flux of 2.5 mL.min^-1^ enabled the time-restricted application of 50 μM nicotine on top of the brain. A baseline recording was performed for 1 min, after which the saline supply was switched to drug supply. The solution reached the *in vivo* preparation within 30 s. The stimulation was maintained for 30 s before switching back to the saline perfusion for an additional 5 min.

Glucose consumption experiments were performed as previously described^21^. Validamycin A (Sigma 32347), a trehalase-selective inhibitor, was directly diluted into artificial hemolymph solution at a final concentration of 40 mM, aliquoted, and stored at −20°C. A freshly thawed aliquot was used for every fly. After 1 min of baseline acquisition, 10 μL of the solution was added to the 90-μL saline droplet on top of the brain, bringing validamycin A to a final concentration of 4 mM. The signal was then acquired for another 12 min. Data were collected indiscriminately from 30 min to 2 h after 5x spaced training. Image analysis was performed using a custom-written MATLAB script. ROI were delimited by hand around the labelled regions of interest (cortex glia or MB neuron somata). The average intensity of the YFP and CFP channels over each ROI was calculated over time after background subtraction. The FRET ratio (YFP/CFP) of the FLII12Pglu-700μδ6 glucose sensor was computed to obtain a signal positively correlated with the glucose concentration. This ratio was normalized by a baseline value calculated over the 30 s preceding drug injection.

For the glucose consumption experiments, the area over the curve (AOC) was computed to obtain values positively correlated with glucose consumption. AOC was calculated as the integral between 200 s and 900 s.

The indicated ‘n’ is the number of animals that were assayed in each condition.

### Generation of transgenic flies

The 2545_pcDNA3.1(-)Pyronic plasmid^30^ was digested by BamHI and BclI. The resulting 2,287 bp fragment was purified by electrophoresis and cloned into a pJFRC19 plasmid (13XLexAop2-IVS-myr::GFP) previously digested by BglII and XbaI to remove the myr::GFP sequence. The resulting construct was verified by restriction. Transgenic fly strains were obtained by site-specific embryonic injection of the resulting vector in the attP18 landing site (chromosome 1), which was outsourced to Rainbow Transgenic Flies, Inc (CA, USA).

### Quantification and statistical analysis

All data are presented as mean ± SEM. For behavior experiments, 2 groups of approximately 30 flies were reciprocally conditioned using either octanol or methylcyclohexanol as the CS^+^. The memory score was calculated from the performance of two groups as described above, which represents an experimental replicate. For imaging experiments, one replicate corresponds to one fly brain. Comparisons of the data series between two conditions were achieved by a two-tailed unpaired t-test. Comparisons between more than two distinct groups were made using a one-way ANOVA test, followed by Tukey’s method for pairwise comparisons between the experimental groups and their controls. ANOVA results are presented as the value of the Fisher distribution F(x,y) obtained from the data, where x is the number of degrees of freedom between groups and y is the total number of degrees of freedom for the distribution. Statistical tests were performed using the GraphPad Prism 8 software. In the figures, asterisks illustrate either the significance level of the t-test or of the least significant pairwise comparison following an ANOVA, with the following nomenclature: *p < 0.05; **p < 0.01; ***p < 0.001; ns: not significant, p > 0.05.

## Acknowledgments

This work was funded by the European Research Council (EnergyMemo Advanced Grant # 741550 to T.P.).

## Author contributions

Conceptualization: P.-Y.P., T.P., and Y.R.; investigation: Y.R. and R.F.; writing – original draft: Y.R. writing – review and editing: P.-Y.P. and T.P.; supervision: P.-Y.P. and T.P.; funding acquisition: T.P.

## Data availability

No datasets that require mandatory deposition into a public database were generated during the current study. Data reported in the current study will be shared by the lead contact upon reasonable request.

## Code availability

This paper does not report original code. Any additional information required to reanalyze the data reported in this paper is available from the lead contact upon request.

