## Supplemental data for "Glycolysis-derived alanine from glia fuels neuronal mitochondria for memory in Drosophila"

### Extended Data Figure Legends

#### Extended Data Figure 1 (related to Figure 1)

a. Without induction of Mpc1 RNAi, memory after single-cycle (1x) training (n = 12, $F_{2,33} = 0.28$, P = 0.76) and spaced training (n = 12, $F_{2,33} = 1.53$, P = 0.23) was normal. b. The behavioral experiments were performed with a second RNAi against Mpc1. Mpc1 knockdown (KD) in adult MB neurons impaired memory after single-cycle training (n = 12, $F_{2,33} = 9.74$, P <0.001) and spaced training (n = 15, $F_{2,42} = 26.03$, P < 0.001), but did not affect memory after single-cycle training followed by cold shock (n = 12, $F_{2,33} = 1.14$, P = 0.33) or massed training (n = 12, $F_{2,33} = 0.14$, P = 0.87). Without induction of Mpc1 RNAi, memory after single-cycle training (n = 10, $F_{2,27} = 0.09$, P = 0.91) and spaced training (n = 11, $F_{2,30} = 0.37$, P = 0.69) was normal. c. Without induction of PDHE1β RNAi, memory after single-cycle training (n = 10, $F_{2,27} = 2.12$, P = 0.14) and spaced training (n = 11-12, $F_{2,31} = 0.60$, P = 0.55) was normal. d. The behavioral experiments were performed with a second RNAi against PDHE1β. PDHE1β knockdown in adult MB neurons impaired memory after single-cycle training (n = 21, $F_{2,60} = 7.05$, P =0.002), but did not affect memory after single-cycle training followed by cold shock (n = 21, $F_{2,60} = 0.48$, P = 0.62). Without induction of PDHE1β RNAi, memory after single-cycle training (n = 12, $F_{2,33} = 0.02$, P = 0.98) was normal. All data are presented as mean ± SEM. Asterisks illustrate the significance level of the t-test, or of the least significant pairwise comparison following an ANOVA, with the following nomenclature: **p < 0.01; ***p < 0.001; ns: not significant, p > 0.05.

#### Extended Data Figure 2 (related to Figure 2)

a. LDH knockdown in adult MB neurons did not affect memory after spaced training (n= 12, $F_{2,33} = 1.73$, P = 0.19). b. Without induction of ALAT RNAi in MB neurons, memory after single-cycle training (n = 12, $F_{2,33} = 0.92$, P = 0.41) and spaced training (n = 12, $F_{2,33} = 0.32$, P = 0.73) was normal. c. The behavioral experiments were performed with a second RNAi against ALAT. ALAT knockdown in adult MB neurons impaired memory after single-cycle training (n = 10, $F_{2,27} = 12.61$, P < 0.001) and spaced training (n =

11, $F_{2,30} = 8.76$, $P = 0.001$), but did not affect memory after single-cycle training followed by cold shock (n = 10, $F_{2,27} = 0.09$, $P = 0.91$) or massed training (n = 10, $F_{2,27} = 1.90$, $P = 0.17$). Without induction of ALAT RNAi in MB neurons, memory after single-cycle training (n = 12, $F_{2,33} = 0.19$, $P = 0.83$) and spaced training (n = 10, $F_{2,27} = 0.77$, $P = 0.47$) was normal. All data are presented as mean ± SEM. Asterisks illustrate the significance level of the t-test, or of the least significant pairwise comparison following an ANOVA, with the following nomenclature: *$p < 0.05$; **$p < 0.01$; ns: not significant, $p > 0.05$.

**Extended Data Figure 3 (related to Figure 3)**

a. Without induction of ALAT RNAi in all glial cells, memory after single-cycle training (n = 10, $F_{2,27} = 0.77$, $P = 0.47$) was normal. b. Without induction of ALAT RNAi in cortex glia, memory after single-cycle training (n = 10, $F_{2,27} = 1.57$, $P = 0.23$) and spaced training (n = 10, $F_{2,27} = 2.51$, $P = 0.10$) was normal. c. The behavioral experiments were performed with a second RNAi against ALAT. ALAT knockdown (KD) in adult cortex glia impaired memory after single-cycle training (n = 10, $F_{2,27} = 5.46$, $P = 0.01$) and spaced training (n = 12, $F_{2,33} = 5.60$, $P = 0.008$), but did not affect memory after single-cycle training followed by cold shock (n = 10, $F_{2,27} = 0.22$, $P = 0.80$) or massed training (n = 12, $F_{2,33} = 0.10$, $P = 0.90$). Without induction of ALAT RNAi in cortex glia, memory after single-cycle training (n = 10, $F_{2,27} = 0.56$, $P = 0.58$) and spaced training (n = 10, $F_{2,27} = 2.34$, $P = 0.12$) was normal. d. The pyruvate sensor Pyronic was expressed in adult MB neurons. The dashed lines delimit the somata of MB neurons where the pyruvate FRET signal was quantified. Scale bar: 50 µm. e. Single-cycle training elicited a faster pyruvate accumulation in MB neuron somata following sodium azide application (5 mM) as compared to non-associative unpaired training (n = 12, $t_{22} = 2.70$, $P = 0.01$). ALAT knockdown in adult cortex glia impaired the single-cycle induced increase in pyruvate accumulation in MB neuron somata following sodium azide application (n = 12, $t_{22} = 0.27$, $P = 0.79$). All data are presented as mean ± SEM. Asterisks illustrate the significance level of the t-test, or of the least significant pairwise comparison following an ANOVA, with the following nomenclature: *$p < 0.05$; ns: not significant, $p > 0.05$.

**Extended Data Figure 4 (related to Figure 4)**

a. Without induction of PFK RNAi in cortex glia, memory after single-cycle training (n = 12, $F_{2,33} = 0.18$, P = 0.84) and spaced training (n = 10, $F_{2,27} = 0.24$, P = 0.79) was normal. b. The behavioral experiments were performed with a second RNAi against PFK. PFK knockdown (KD) in adult cortex glia had no significant effect on total memory after single-cycle training (n = 13, $F_{2,36} = 1.60$, P = 0.22) or memory after single-cycle training followed by cold shock (n = 13, $F_{2,36} = 4.56$, P = 0.02). Instead, there was a non-significant trend towards a decreased total 3h-memory and increased memory after cold shock (MT-ARM component), which can mask potential effects on MTM. By subtracting the residual memory after cold shock from the total memory expressed 3 h after single-cycle training, we were able to display only the MTM. PFK knockdown in adult cortex glia impaired MTM after single-cycle training (n = 13, $F_{2,36} = 5.23$, P = 0.01) and spaced training (n = 20, $F_{2,57} = 5.10$, P = 0.009), but had no effect on massed training (n = 14, $F_{2,39} = 3.05$, P = 0.06). Without induction of PFK RNAi in cortex glia, memory after single-cycle training (n = 12, $F_{2,33} = 0.60$, P = 0.55) and single-cycle training followed by cold shock (n = 12, $F_{2,33} = 0.41$, P = 0.67) were normal. Without induction of PFK RNAi, MTM (n = 12, $F_{2,33} = 0.38$, P = 0.69) and memory after spaced training (n = 18, $F_{2,51} = 0.35$, P = 0.70) were normal. c. Single-cycle training elicited a faster pyruvate accumulation in MB neuron somata following sodium azide application (5 mM) as compared to non-associative unpaired training (n = 12, $t_{22} = 3.36$, P = 0.003). PFK knockdown in adult cortex glia impaired the single-cycle induced increase in pyruvate accumulation in MB neuron somata following sodium azide application (n = 12, $t_{22} = 1.88$, P = 0.07). All data are presented as mean ± SEM. Asterisks illustrate the significance level of the t-test, or of the least significant pairwise comparison following an ANOVA, with the following nomenclature: *p < 0.05; **p < 0.01; ns: not significant, p > 0.05.

**Extended Data Figure 5 (related to Figure 5)**

a. Treh knockdown in adult cortex glia did not affect memory after single-cycle training (n = 10, $F_{2,27} = 0.20$, P = 0.82) or single-cycle training followed by cold shock (n = 10, $F_{2,27} = 0.69$, P = 0.51). b. GlyP knockdown in adult cortex glia did not affect memory after single-cycle training (n = 14, $F_{2,36} = 0.64$, P = 0.53) or single-cycle training followed by cold shock (n = 14, $F_{2,36} = 0.41$, P = 0.67). c. Transcriptomic data for the family of SLC2 hexose sugar transporters expressed in cortex glia from[35]. The SLC2 family is composed of 31 putative hexose sugar transporters. Nebu is the most abundant member in the cortex glia, followed by glug. d. Without induction of glug RNAi in cortex glia, memory after single-cycle training (n = 12, $F_{2,33} = 0.40$, P = 0.67) and spaced training (n = 10, $F_{2,27} = 0.23$, P = 0.79) was normal. e. Single-cycle training elicited a faster pyruvate accumulation in MB neuron somata following sodium azide application (5 mM) as compared to non-associative unpaired training (n = 12, $t_{22} = 3.89$, P = 0.0008). Glug knockdown in adult cortex glia impaired the single-cycle induced increase in pyruvate accumulation in MB neuron somata following sodium azide application (n = 12, $t_{22} = 0.66$, P = 0.62). Nebu knockdown in adult cortex glia did not impair the single-cycle induced increase in pyruvate accumulation in MB neuron somata following sodium azide application (n = 12, $t_{22} = 3.21$, P = 0.004). All data are presented as mean ± SEM. Asterisks illustrate the significance level of the t-test, or of the least significant pairwise comparison following an ANOVA, with the following nomenclature: **p < 0.01; ***p < 0.001; ns: not significant, p > 0.05.

**Extended Data Figure 6 (related to Figure 6)**

a. Without induction of nebu RNAi in cortex glia, memory after spaced training (n = 10, $F_{2,27} = 1.31$, P = 0.29) was normal. b. The behavioral experiments were performed with a second RNAi against nebu. Nebu knockdown (KD) in adult cortex glia impaired spaced training (n = 12, $F_{2,33} = 7.47$, P = 0.002), but did not affect memory after single-cycle training (n = 15, $F_{2,42} = 2.40$, P = 0.10), single-cycle training followed by cold shock (n = 15, $F_{2,42} = 1.15$, P = 0.33), or massed training (n = 10, $F_{2,27} = 0.96$, P = 0.40). Without induction of nebu RNAi, memory after spaced training (n = 10, $F_{2,27} = 0.60$, P = 0.55) was normal. All data are presented as mean ± SEM. Asterisks illustrate the significance level of the t-test, or of the least significant pairwise comparison following an ANOVA, with the following nomenclature: *p < 0.05; ns: not significant, p > 0.05.

**Extended Data Tables**

Extended Data Table 1. Sensory acuity controls related to Figure 1 and Extended Data Figure 1.

| Genotypes | Shock reactivity | | Olfactory acuity | | | |
|---|---|---|---|---|---|---|
| | | | Octanol | | Methylcyclohexanol | |
| | Mean ± SEM | Statistics | Mean ± SEM | Statistics | Mean ± SEM | Statistics |
| tub-GAL80$^{ts}$; VT30559-GAL4/+ | 54.5 ± 4.8 | n = 12 | 57.1 ± 4.9 | n = 12 | 52.8 ± 4.9 | n = 12 |
| +/UAS-Mpc1 RNAi HMS05634 | 49.5 ± 3.8 | $F_{2,33}$ = 0.31 | 47.9 ± 4.7 | $F_{2,33}$= 1.02 | 49.6 ± 3.1 | $F_{2,33}$ = 0.76 |
| tub-GAL80$^{ts}$; VT30559-GAL4> UAS-Mpc1 RNAi HMS05634 | 53.0 ± 5.2 | P = 0.74 | 55.8 ± 5.2 | P = 0.37 | 52.3 ± 4.1 | P = 0.48 |
| tub-GAL80$^{ts}$; VT30559-GAL4/+ | 68.1 ± 5.2 | n = 12 | 50.6 ± 3.2 | n = 12 | 46.0 ± 5.7 | n = 12 |
| +/UAS-PDHE1β RNAi HMC03762 | 59.8 ± 4.9 | $F_{2,33}$ = 1.61 | 50.0 ± 6.3 | $F_{2,33}$ = 2.70 | 44.6 ± 5.3 | $F_{2,33}$ = 0.03 |
| tub-GAL80$^{ts}$; VT30559-GAL4> UAS-PDHE1β RNAi HMC03762 | 71.6 ± 4.2 | P = 0.22 | 63.8 ± 4.2 | P = 0.08 | 46.5 ± 6.0 | P = 0.97 |
| tub-GAL80$^{ts}$; VT30559-GAL4/+ | 55.2 ± 3.7 | n = 12 | 56.2 ± 3.5 | n = 10 | 55.2 ± 5.4 | n = 10 |
| +/UAS-Mpc1 RNAi KK102734 | 54.6 ± 5.3 | $F_{2,33}$ = 0.01 | 48.2 ± 4.4 | $F_{2,27}$ = 1.74 | 42.9 ± 6.6 | $F_{2,27}$ = 2.22 |
| tub-GAL80$^{ts}$; VT30559-GAL4> UAS-Mpc1 RNAi KK102734 | 54.6 ± 6.5 | P > 0.99 | 56.2 ± 2.2 | P = 0.19 | 58.7 ± 5.0 | P = 0.14 |

Extended Data Table 2. Sensory acuity controls related to Figure 2 and Extended Data Figure 2.

<table>
<tr><th rowspan="3">Genotypes</th><th colspan="2" rowspan="2">Shock reactivity</th><th colspan="4">Olfactory acuity</th></tr>
<tr><th colspan="2">Octanol</th><th colspan="2">Methylcyclohexanol</th></tr>
<tr><th>Mean ± SEM</th><th>Statistics</th><th>Mean ± SEM</th><th>Statistics</th><th>Mean ± SEM</th><th>Statistics</th></tr>
<tr><td>tub-GAL80$^{ts}$; VT30559-GAL4/+</td><td>53.4 ± 3.6</td><td rowspan="3">n = 10<br>$F_{2,27}$ = 0.25<br>P = 0.78</td><td>60.2 ± 5.2</td><td rowspan="3">n = 10<br>$F_{2,27}$ = 1.70<br>P = 0.20</td><td>70.1 ± 5.0</td><td rowspan="3">n = 10<br>$F_{2,27}$ = 0.03<br>P = 0.97</td></tr>
<tr><td>+/UAS-ALAT RNAi GD9174</td><td>57.5 ± 5.6</td><td>62.7 ± 3.9</td><td>71.3 ± 5.8</td></tr>
<tr><td>tub-GAL80$^{ts}$; VT30559-GAL4> UAS-ALAT RNAi GD9174</td><td>57.9 ± 5.6</td><td>71.9 ± 5.0</td><td>71.7 ± 4.6</td></tr>
<tr><td>tub-GAL80$^{ts}$; VT30559-GAL4/+</td><td>54.8 ± 5.1</td><td rowspan="3">n = 10<br>$F_{2,27}$ = 0.34<br>P = 0.72</td><td>72.3 ± 4.5</td><td rowspan="3">n = 10<br>$F_{2,27}$ = 0.48<br>P = 0.70</td><td>82.5 ± 3.8</td><td rowspan="3">n = 10<br>$F_{2,27}$ = 1.33<br>P = 0.28</td></tr>
<tr><td>+/UAS-ALAT RNAi HMC05124</td><td>50.1 ± 3.6</td><td>73.7 ± 5.6</td><td>75.9 ± 4.6</td></tr>
<tr><td>tub-GAL80$^{ts}$; VT30559-GAL4> UAS-ALAT RNAi HMC05124</td><td>52.5 ± 3.2</td><td>65.7 ± 5.1</td><td>80.7 ± 5.7</td></tr>
</table>

Extended Data Table 3. Sensory acuity controls related to Figure 3 and Extended Data Figure 3.

| Genotypes | Shock reactivity | | Olfactory acuity | | | |
|---|---|---|---|---|---|---|
| | | | Octanol | | Methylcyclohexanol | |
| | Mean ± SEM | Statistics | Mean ± SEM | Statistics | Mean ± SEM | Statistics |
| tub-GAL80$^{ts}$; 54H02-GAL4/+ | 62.6 ± 7.0 | n = 12 $F_{2,33}$= 0.34 P = 0.72 | 68.1 ± 5.1 | n = 12 $F_{2,33}$ = 0.40 P = 0.67 | 64.9 ± 2.6 | n = 12 $F_{2,33}$= 1.27 P = 0.29 |
| +/UAS-ALAT RNAi GD9174 | 55.3 ± 6.9 | | 70.2 ± 7.1 | | 72.1 ± 5.3 | |
| tub-GAL80$^{ts}$; 54H02-GAL4> UAS-ALAT RNAi GD9174 | 60.9 ± 5.7 | | 75.4 ± 5.3 | | 62.3 ± 5.1 | |
| tub-GAL80$^{ts}$; 54H02-GAL4/+ | 62.5 ± 5.0 | n = 10 $F_{2,27}$ = 0.97 P = 0.39 | 79.4 ± 3.1 | n = 10 $F_{2,27}$ = 1.40 P = 0.26 | 78.7 ± 3.3 | n = 10 $F_{2,27}$ = 1.04 P = 0.39 |
| +/UAS-ALAT RNAi HMC05124 | 55.5 ± 5.9 | | 66.6 ± 4.6 | | 78.5 ± 4.8 | |
| tub-GAL80$^{ts}$; 54H02-GAL4> UAS-ALAT RNAi HMC05124 | 65.4 ± 4.6 | | 73.3 ± 5.4 | | 80.0 ± 4.6 | |

Extended Data Table 4. Sensory acuity controls related to Figure 4 and Extended Data Figure 4.

| Genotypes | Shock reactivity | | Olfactory acuity | | | |
|---|---|---|---|---|---|---|
| | | | Octanol | | Methylcyclohexanol | |
| | Mean ± SEM | Statistics | Mean ± SEM | Statistics | Mean ± SEM | Statistics |
| tub-GAL80ts; 54H02-GAL4/+ | 51.7 ± 3.3 | n = 10 $F_{2,27}$ = 1.24 P = 0.30 | 60.4 ± 4.7 | n = 10 $F_{2,27}$ =1.95 P = 0.14 | 65.5 ± 4.6 | n = 10 $F_{2,27}$ = 0.63 P = 0.60 |
| +/UAS-PFK RNAi KK101887 | 59.0 ± 6.0 | | 73.6 ± 4.7 | | 60.1 ± 5.7 | |
| tub-GAL80ts; 54H02-GAL4> UAS-PFK RNAi KK101887 | 63.1 ± 5.8 | | 60.2 ± 5.1 | | 66.8 ± 5.7 | |
| tub-GAL80ts; 54H02-GAL4/+ | 66.7 ± 3.4 | n = 10 $F_{2,27}$ = 0.69 P = 0.51 | 72.4 ± 3.7 | n = 10 $F_{2,27}$ = 0.58 P = 0.63 | 72.7 ± 5.6 | n = 10 $F_{2,27}$ = 0.79 P = 0.51 |
| +/UAS-PFK RNAi HMS01324 | 64.8 ± 3.8 | | 74.2 ± 4.4 | | 66.7 ± 4.1 | |
| tub-GAL80ts; 54H02-GAL4> UAS-PFK RNAi HMS01324 | 60.8 ± 3.6 | | 66.1 ± 5.1 | | 77.1 ± 4.9 | |

Extended Data Table 5. Sensory acuity controls related to Figure 5 and Extended Data Figure 5.

| Genotypes | Shock reactivity | | Olfactory acuity | | | |
|---|---|---|---|---|---|---|
| | | | Octanol | | Methylcyclohexanol | |
| | Mean ± SEM | Statistics | Mean ± SEM | Statistics | Mean ± SEM | Statistics |
| tub-GAL80ts; 54H02-GAL4/+ | 58.9 ± 5.2 | n = 10 $F_{2,27}$ = 0.55 P = 0.65 | 66.1 ± 5.1 | n = 10 $F_{2,27}$ = 0.73 P = 0.54 | 62.9 ± 5.4 | n = 10 $F_{2,27}$ = 0.1 P = 0.91 |
| +/UAS-glug RNAi GD2869 | 65.9 ± 5.5 | | 64.3 ± 4.7 | | 65.9 ± 4.4 | |
| tub-GAL80ts; 54H02-GAL4> UAS-glug RNAi GD2869 | 61.1 ± 5.9 | | 60.2 ± 4.4 | | 65.5 ± 5.8 | |

Extended Data Table 6. Sensory acuity controls related to Figure 6 and Extended Data Figure 6.

| Genotypes | Shock reactivity | | Olfactory acuity | | | |
|---|---|---|---|---|---|---|
| | | | Octanol | | Methylcyclohexanol | |
| | Mean ± SEM | Statistics | Mean ± SEM | Statistics | Mean ± SEM | Statistics |
| tub-GAL80$^{ts}$; 54H02-GAL4/+ | 50.9 ± 3.0 | n = 10<br>$F_{2,27}$ = 1.34<br>P = 0.28 | 73.1 ± 3.7 | n = 10<br>$F_{2,27}$ = 0.32<br>P = 0.73 | 77.6 ± 3.9 | n = 10<br>$F_{2,27}$ = 0.25<br>P = 0.78 |
| +/UAS-nebu RNAi HMS01072 | 52.4 ± 4.7 | | 77.2 ± 3.6 | | 75.1 ± 3.7 | |
| tub-GAL80$^{ts}$; 54H02-GAL4> UAS-nebu RNAi HMS01072 | 52.5 ± 4.6 | | 77.5 ± 5.5 | | 79.3 ± 5.1 | |
| tub-GAL80$^{ts}$; 54H02-GAL4/+ | 62.0 ± 4.0 | n = 10<br>$F_{2,27}$ = 0.06<br>P = 0.95 | 65.9 ± 4.4 | n = 10<br>$F_{2,27}$ = 2.02<br>P = 0.15 | 72.4 ± 4.1 | n = 10<br>$F_{2,27}$ = 0.06<br>P = 0.95 |
| +/UAS-nebu RNAi GD2444 | 63.5 ± 3.2 | | 72.2 ± 4.1 | | 70.7 ± 4.8 | |
| tub-GAL80$^{ts}$; 54H02-GAL4> UAS-nebu RNAi GD2444 | 62.2 ± 3.4 | | 61.1 ±3.1 | | 72.6 ± 4.9 | |

Extended Data Table 7. Drosophila strains used in this study.

| **Fly strain** | **Stock n°/Source/ reference** |
|---|---|
| VT30559-GAL4 | VDRC 206077 |
| repo-GAL4 | BDSC 7415 |
| alrm-GAL4 | [61] |
| 56F03-GAL4 | BDSC 39157 |
| 54H02-GAL4 | [62] |
| 13F02-LexA | BDSC 52460 |
| tub-GAL80$^{ts}$ | BDSC 7019 |
| tub-GAL80$^{ts}$; VT30559-GAL4 | [23] |
| tub-GAL80$^{ts}$; repo-GAL4 | This report |
| tub-GAL80$^{ts}$; alrm-GAL4 | [21] |
| tub-GAL80$^{ts}$; 56F03-GAL4 | [21] |
| tub-GAL80$^{ts}$; 54H02-GAL4 | [21] |
| tub-GAL80$^{ts}$; VT30559-GAL4, UAS-Pyronic | [23] |
| tub-GAL80$^{ts}$, 13F02-LexA; 54H02-GAL4 | [21] |
| tub-GAL80$^{ts}$; 54H02-GAL4, UAS-FLII12Pglu-700μδ6 | [21] |
| UAS-Mpc1 RNAi HMS05634 | BDSC 67817 |
| UAS-Mpc1 RNAi KK102734 | VDRC 103829 |
| UAS-PDHE1β RNAi HMC03762 | BDSC 55619 |
| UAS-PDHE1β RNAi KK107865 | VDRC 104022 |
| UAS-PFK RNAi KK101887 | VDRC 105666 |
| UAS-PFK RNAi HMS01324 | BDSC 34336 |
| UAS-LDH RNAi KK102330 | VDRC 110190 |
| UAS-ALAT RNAi GD9174 | VDRC 32681 |
| UAS-ALAT RNAi HMC05124 | BDSC 60130 |
| UAS-glug RNAi GD2869 | VDRC 42627 |
| UAS-nebu RNAi HMS01072 | BDSC 34598 |

| | |
|---|---|
| UAS-nebu RNAi GD2444 | VDRC 8359 |
| UAS-Treh RNAi HMC03381 | BDSC 51810 |
| UAS-GlyP RNAi GD12183 | VDRC 27928 |
| UAS-Pyronic | [23] |
| UAS-FLII12Pglu-700μδ6 | [58] |
| LexAop-FLII12Pglu-700μδ6 | [21] |
| LexAop-FLII12Pglu-700μδ6; UAS-glug RNAi GD2869 | This report |
| LexAop-FLII12Pglu-700μδ6; UAS-nebu RNAi HMS01072 | This report |
| LexAop-Pyronic; UAS-ALAT RNAi GD9174 | This report |
| LexAop-Pyronic; UAS-PFK RNAi HMS01324 | This report |
| LexAop-Pyronic; UAS-glug RNAi GD2869 | This report |
| LexAop-Pyronic; UAS-nebu RNAi HMS01072 | This report |

Extended Data Figure 1

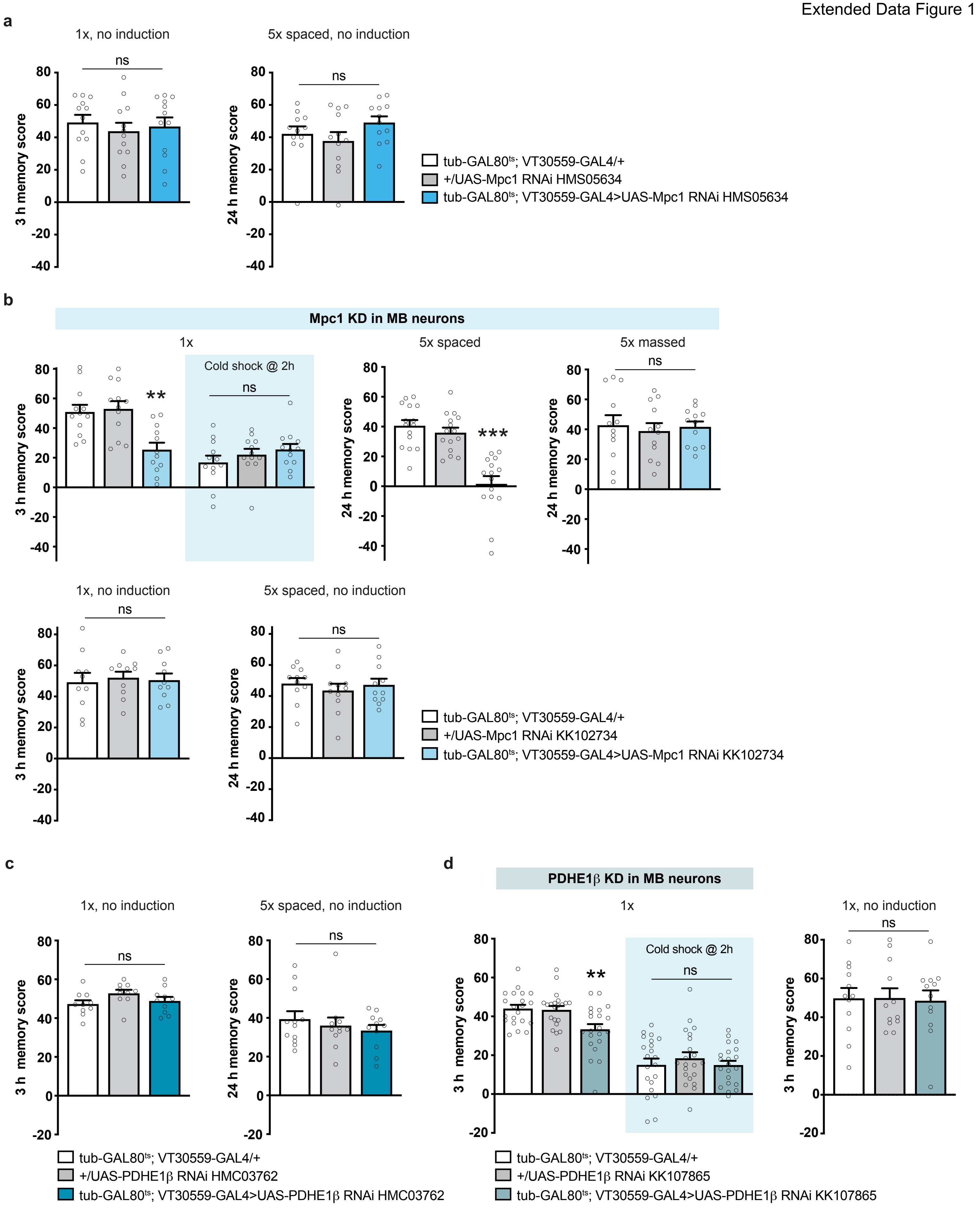

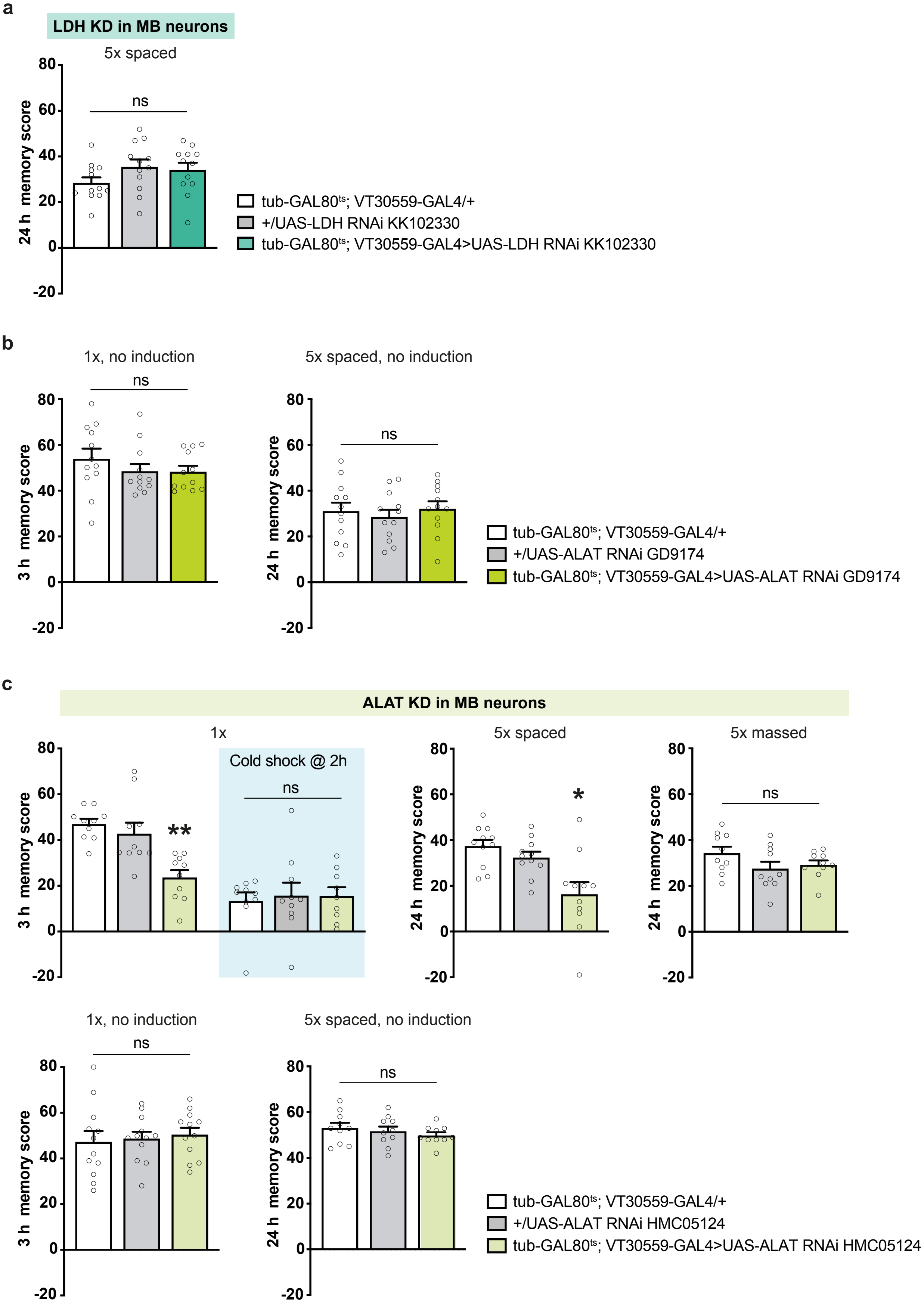
a
LDH KD in MB neurons
5x spaced
ns
24 h memory score
tub-GAL80ts; VT30559-GAL4/+
+/UAS-LDH RNAi KK102330
tub-GAL80ts; VT30559-GAL4>UAS-LDH RNAi KK102330
b
1x, no induction
5x spaced, no induction
3 h memory score
24 h memory score
tub-GAL80ts; VT30559-GAL4/+
+/UAS-ALAT RNAi GD9174
tub-GAL80ts; VT30559-GAL4>UAS-ALAT RNAi GD9174
c
ALAT KD in MB neurons
1x
Cold shock @ 2h
5x spaced
5x massed
**
*
1x, no induction
5x spaced, no induction
tub-GAL80ts; VT30559-GAL4/+
+/UAS-ALAT RNAi HMC05124
tub-GAL80ts; VT30559-GAL4>UAS-ALAT RNAi HMC05124

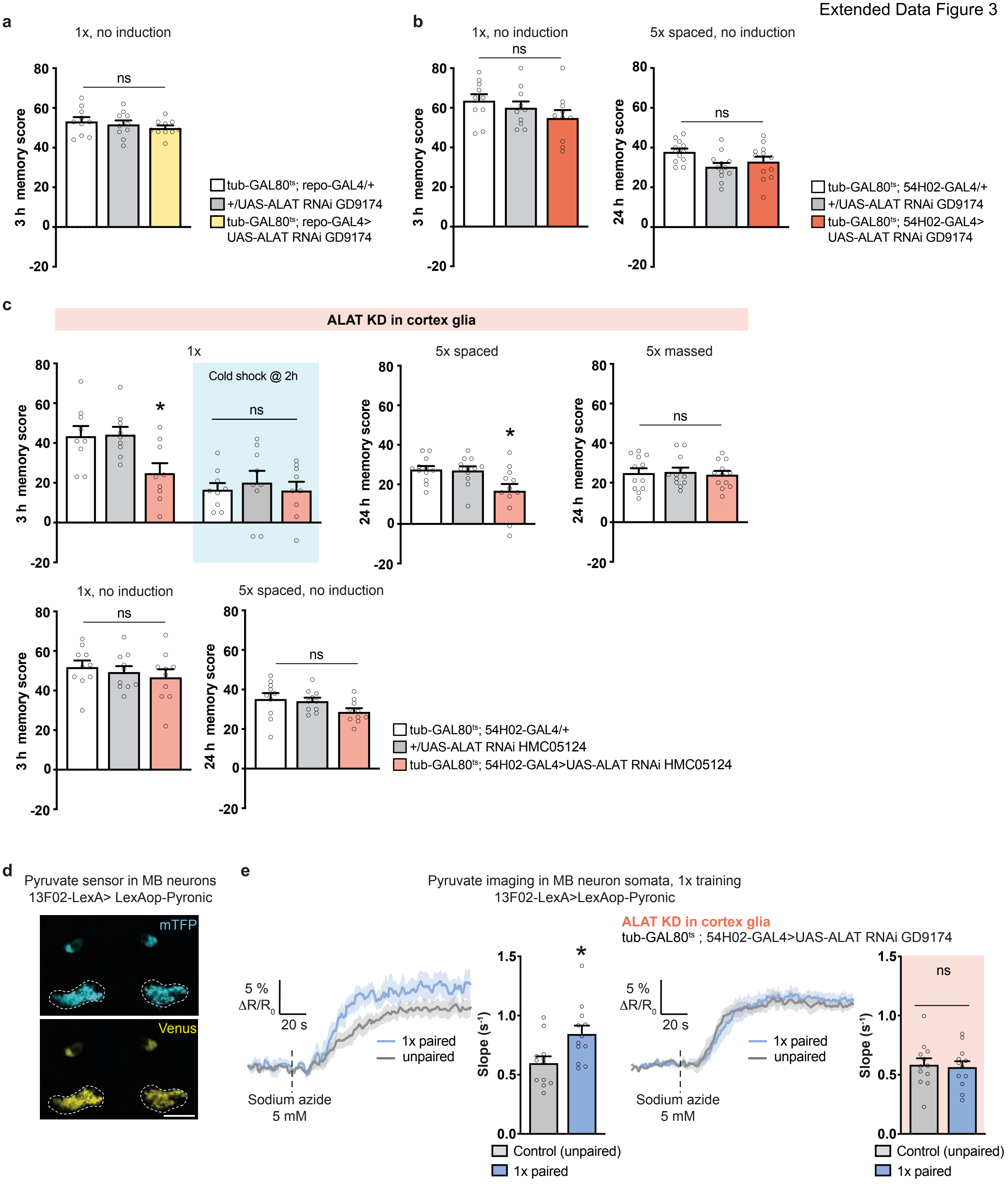
a
1x, no induction
ns
3 h memory score
tub-GAL80ts; repo-GAL4/+
+/UAS-ALAT RNAi GD9174
tub-GAL80ts; repo-GAL4>
UAS-ALAT RNAi GD9174
b
1x, no induction
ns
5x spaced, no induction
ns
24 h memory score
tub-GAL80ts; 54H02-GAL4/+
+/UAS-ALAT RNAi GD9174
tub-GAL80ts; 54H02-GAL4>
UAS-ALAT RNAi GD9174
c
ALAT KD in cortex glia
1x
Cold shock @ 2h
5x spaced
5x massed
1x, no induction
5x spaced, no induction
tub-GAL80ts; 54H02-GAL4/+
+/UAS-ALAT RNAi HMC05124
tub-GAL80ts; 54H02-GAL4>UAS-ALAT RNAi HMC05124
d
Pyruvate sensor in MB neurons
13F02-LexA> LexAop-Pyronic
mTFP
Venus
e
Pyruvate imaging in MB neuron somata, 1x training
13F02-LexA>LexAop-Pyronic
ALAT KD in cortex glia
tub-GAL80ts ; 54H02-GAL4>UAS-ALAT RNAi GD9174
5 %
ΔR/R0
20 s
1x paired
unpaired
Sodium azide
5 mM
Slope (s-1)
Control (unpaired)
1x paired
ns

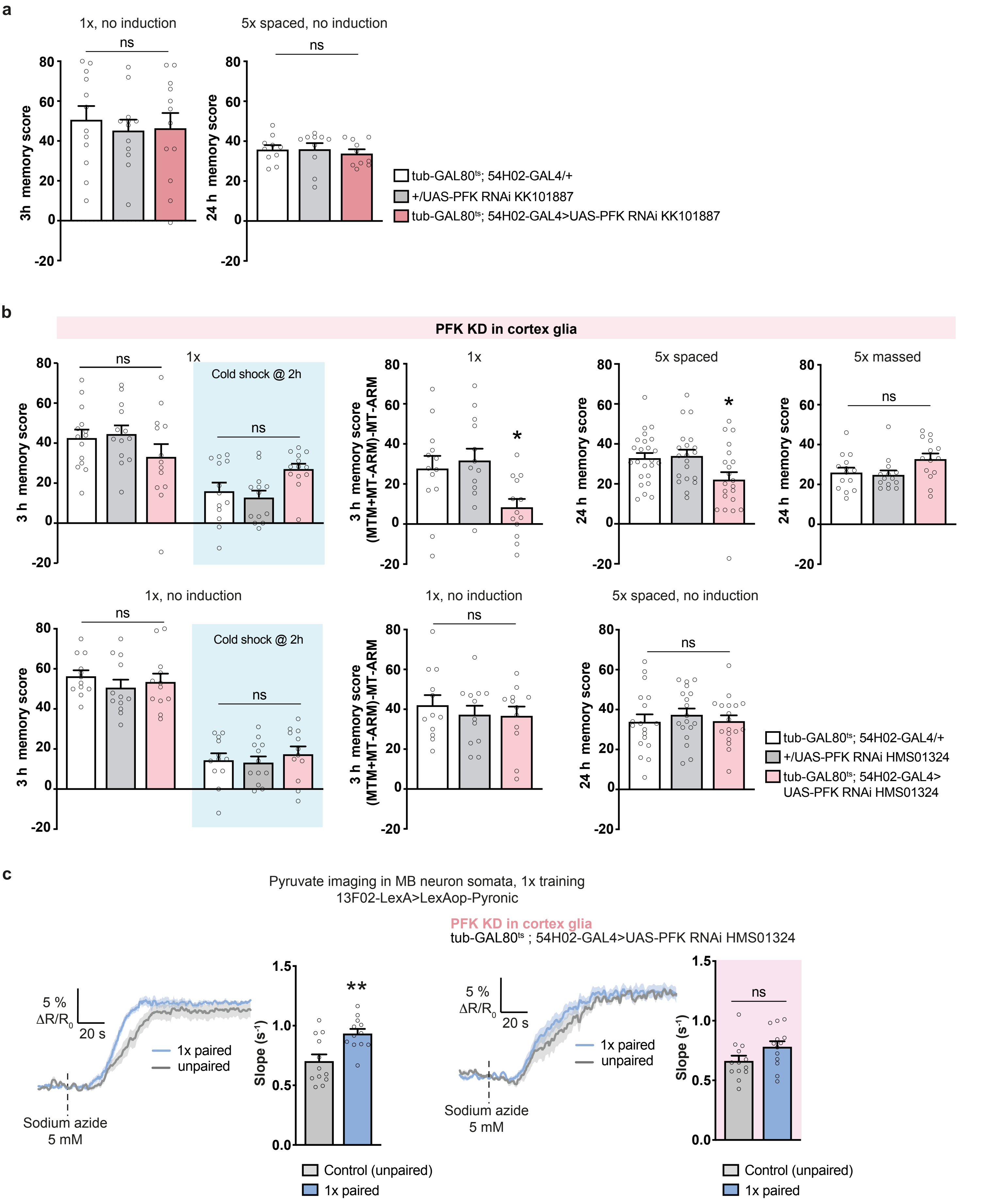
a
1x, no induction
ns
3h memory score
5x spaced, no induction
ns
24 h memory score
tub-GAL80ts; 54H02-GAL4/+
+/UAS-PFK RNAi KK101887
tub-GAL80ts; 54H02-GAL4>UAS-PFK RNAi KK101887
b
PFK KD in cortex glia
1x
Cold shock @ 2h
ns
3 h memory score
3 h memory score (MTM+MT-ARM)-MT-ARM
5x spaced
24 h memory score
5x massed
1x, no induction
5x spaced, no induction
tub-GAL80ts; 54H02-GAL4/+
+/UAS-PFK RNAi HMS01324
tub-GAL80ts; 54H02-GAL4>UAS-PFK RNAi HMS01324
c
Pyruvate imaging in MB neuron somata, 1x training
13F02-LexA>LexAop-Pyronic
PFK KD in cortex glia
tub-GAL80ts ; 54H02-GAL4>UAS-PFK RNAi HMS01324
5 % ΔR/R0
20 s
1x paired
unpaired
Sodium azide 5 mM
Slope (s-1)
Control (unpaired)
1x paired
ns

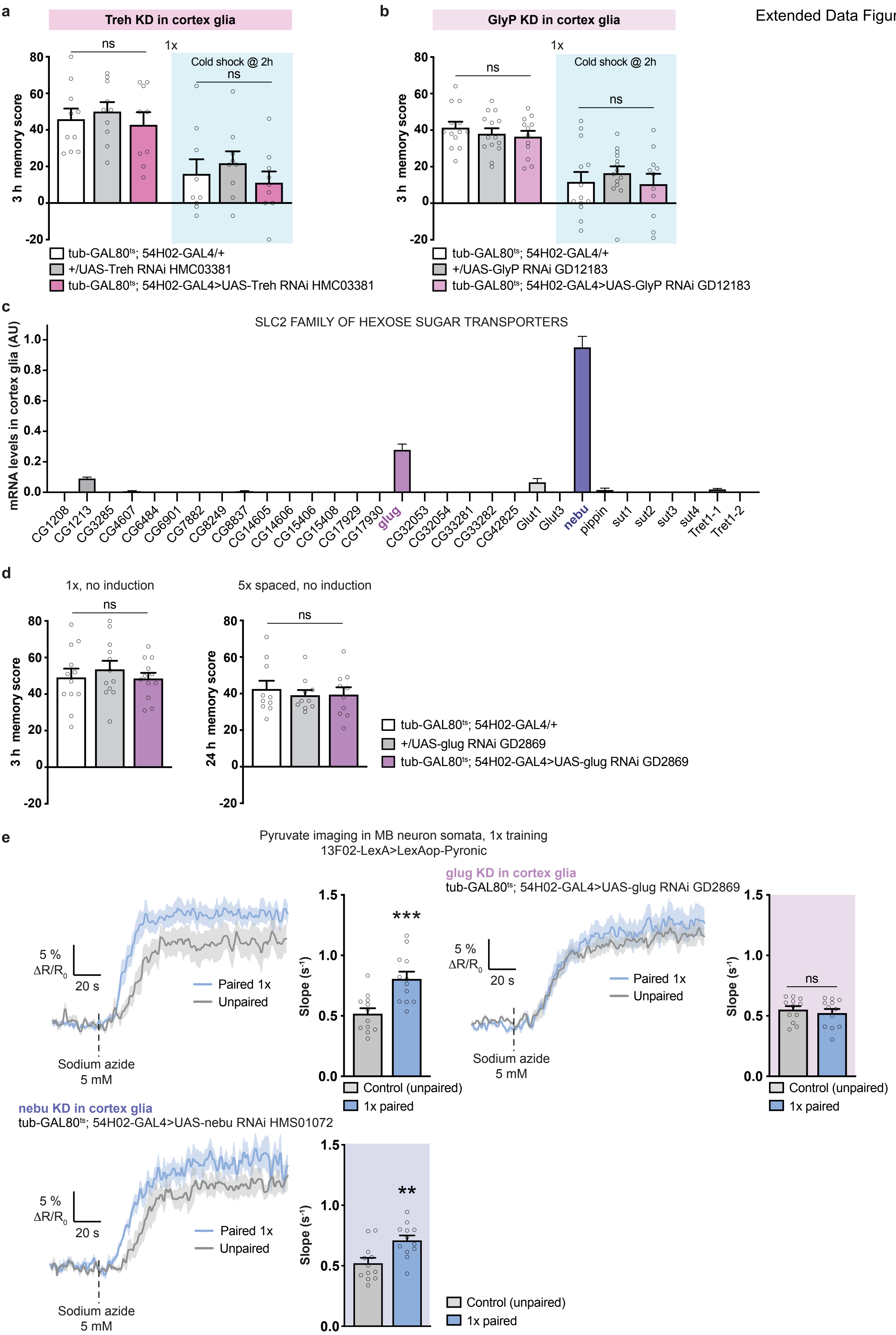
a
Treh KD in cortex glia
1x
Cold shock @ 2h
ns
ns
3 h memory score
tub-GAL80ts; 54H02-GAL4/+
+/UAS-Treh RNAi HMC03381
tub-GAL80ts; 54H02-GAL4>UAS-Treh RNAi HMC03381
b
GlyP KD in cortex glia
1x
Cold shock @ 2h
ns
ns
3 h memory score
tub-GAL80ts; 54H02-GAL4/+
+/UAS-GlyP RNAi GD12183
tub-GAL80ts; 54H02-GAL4>UAS-GlyP RNAi GD12183
c
SLC2 FAMILY OF HEXOSE SUGAR TRANSPORTERS
mRNA levels in cortex glia (AU)
CG1208 CG1213 CG3285 CG4607 CG6484 CG6901 CG7882 CG8249 CG8837 CG14605 CG14606 CG15406 CG15408 CG17929 CG17930 glug CG32053 CG32054 CG33281 CG33282 CG42825 Glut1 Glut3 nebu pippin sut1 sut2 sut3 sut4 Tret1-1 Tret1-2
d
1x, no induction
5x spaced, no induction
ns
ns
3 h memory score
24 h memory score
tub-GAL80ts; 54H02-GAL4/+
+/UAS-glug RNAi GD2869
tub-GAL80ts; 54H02-GAL4>UAS-glug RNAi GD2869
e
Pyruvate imaging in MB neuron somata, 1x training
13F02-LexA>LexAop-Pyronic
5 % ΔR/R0
20 s
Paired 1x
Unpaired
Sodium azide 5 mM
Slope (s-1)
***
Control (unpaired)
1x paired
glug KD in cortex glia
tub-GAL80ts; 54H02-GAL4>UAS-glug RNAi GD2869
ns
nebu KD in cortex glia
tub-GAL80ts; 54H02-GAL4>UAS-nebu RNAi HMS01072
**

Extended Data Figure 6

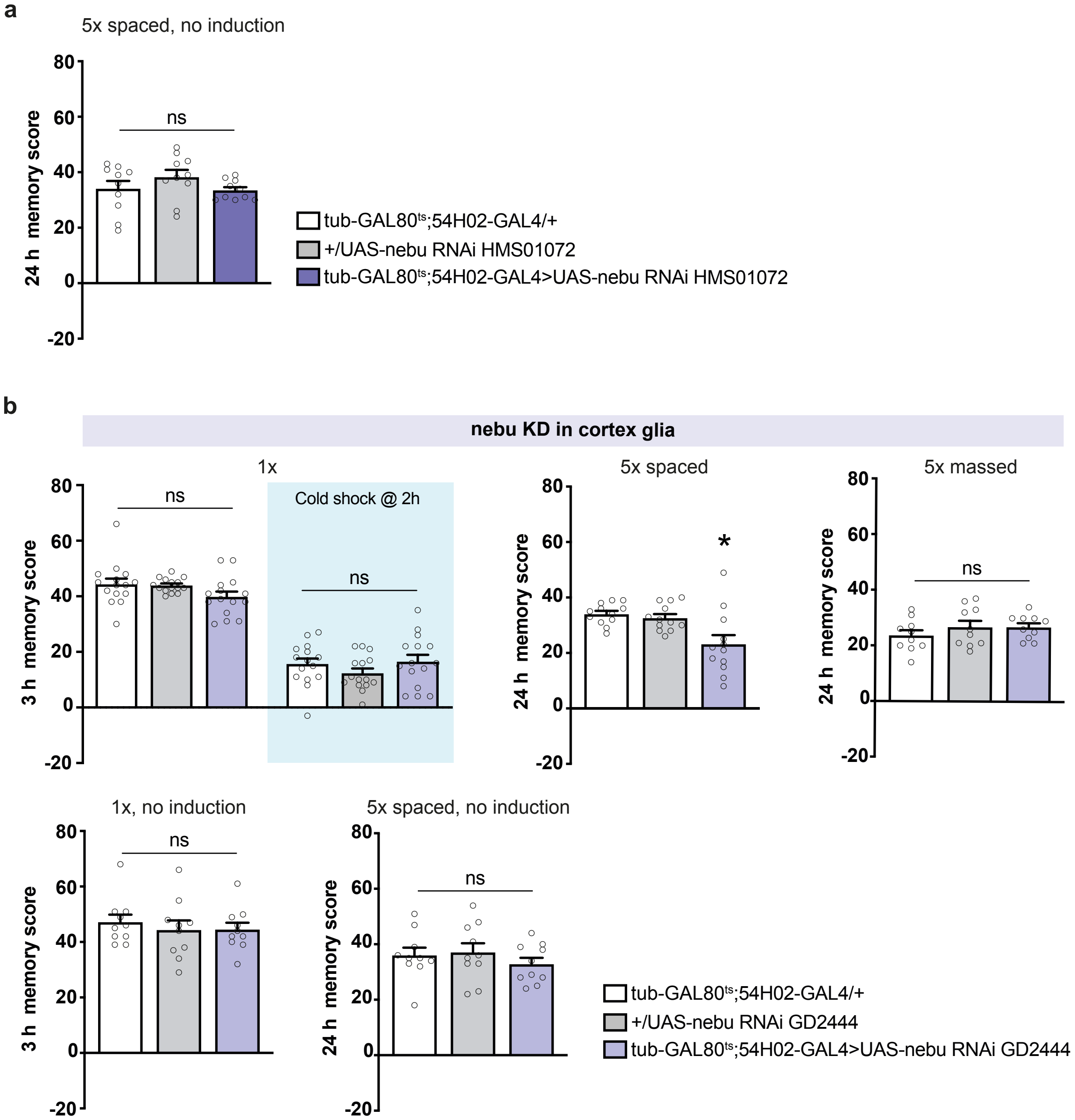
